# The Nature of *Espeletia* Species

**DOI:** 10.1101/2020.09.29.318865

**Authors:** Yam M. Pineda, Andrés J. Cortés, Santiago Madriñán, Iván Jiménez

## Abstract

Species are often regarded as basic units of study in biology, following the presumption that they are real and discrete natural entities. But several biologists wonder if species are arbitrary divisions that do not correspond to discrete natural groups of organisms. Two issues must be addressed to solve this controversy, but few studies seem to do so. The first is whether organisms form sympatric and synchronic groups that are distinct in terms of phenotypes and genome-wide allele frequencies, often called “good species.” Alternatives to “good species” include “cryptic species,” syngameons and, more generally, cases in which phenotypes and genome-wide allele frequencies reflect contrasting evolutionary histories. The second issue is the degree to which species taxa (i.e., taxonomic classification at the species level) reflect natural groups of organisms or constitute arbitrary divisions of biological diversity. Here, we empirically addressed both issues by studying plants of the Andean genus *Espeletia* (Asteraceae). We collected a geographically dense sample of 538 specimens from the paramo de Sumapaz, in the Cordillera Oriental of Colombia. Additionally, we examined 165 herbarium specimens previously collected by other researchers in this region, or from taxa known to occur there. We tested for the existence of phenotypic groups using normal mixture models and data on 13 quantitative characters. Among 307 specimens with all 13 measurements, we found six distinct phenotypic groups in sympatry. We also tested for the existence of groups defined by genome-wide allele frequencies, using ancestry models and data on 2,098 single nucleotide polymorphisms. Among 77 specimens with complete genomic data, we found three groups in sympatry, with high levels of admixture. Concordance between groups defined by phenotype and genome-wide allele frequencies was low, suggesting that phenotypes and genome-wide allele frequencies reflect contrasting evolutionary histories. Moreover, the high levels of admixture suggest that *Espeletia* plants form a syngameon in the paramo de Sumapaz. To determine the extent to which species taxa corresponded to phenotypic and genomic groups, we used data on 12 phenotypic characters to assign 307 specimens to species taxa, according to descriptions of species taxa in the most recent monograph of *Espeletia*. This sample included 27 specimens cited in the monograph. Remarkably, only one out of 307 specimens in our sample fell inside any of the phenotypic ranges reported in the monograph for the species taxa known to occur in the paramo de Sumapaz. These results show that species taxa in *Espeletia* are delineations of largely empty phenotypic space that miss biological diversity.

---

Species are often regarded as basic units of biological diversity in ecology, evolution, biogeography and conservation biology (Richards 2010; Sigwart 2018), following the presumption that they are real, discrete natural entities (Coyne and Orr 2004; Barraclough and Humphreys 2015). But some biologists, most notoriously botanists, have persistent doubts about the existence of species. They see species as arbitrary divisions of biological diversity that do not necessarily correspond to discrete natural groups of organisms (Levin 1979; Raven 1986; Bachmann 1998). Darwin appeared to have developed a similar view (Mayr 1982; Stamos 2007; Richards 2010; Mallet 2013 but see De Queiroz 2011; Wilkins 2018), seemingly revealed in the concluding chapter of On the Origin of Species: ‘‘…we shall have to treat species in the same manner as those naturalists treat genera, who admit that genera are merely artificial combinations made for convenience. This may not be a cheering prospect, but we shall at least be freed from the vain search for the undiscovered and undiscoverable essence of the term species’’ (Darwin 1859). Moreover, it has been argued that Darwin’s view on the nature of species may have been shaped by interactions with botanists (Mayr 1982), and that botanists could have been overly impressed by a few examples of fuzzy species limits (e.g., oaks, hawthorns and blackberries) and wrongly assumed these “botanical horror stories” to be representative of plant species overall (Diamond 1992). Yet, skepticism about the nature of species is still common among botanists (Barraclough and Humphreys 2015; Hipp et al. 2019) and other biologists (e.g., Hey 2001), as this fundamental question about the structure of biodiversity, with major basic and applied implications, remains unsatisfactorily addressed (Barraclough 2019).

Attempts to empirically solve the controversy about the nature of species would ideally address two related but separate issues. The first is whether coeval individual organisms form sympatric, distinct groups in nature (Coyne and Orr 2004). Sympatry, defined in terms of the normal cruising range of individual organisms (Mayr 1947; Mallet et al. 2009), and synchrony are stressed in this context because they imply opportunity for two or more groups to merge via hybridization and introgression (Coyne and Orr 2004). Additionally, temporal and spatial co-occurrence implies opportunity for competitive exclusion. Therefore, distinct groups of organisms are unlikely to coexist for long unless ecological niche differences between them are enough to offset respective differences in population growth rate (Chesson 2000; Adler et al. 2007). In short, when sympatric and synchronous, discrete groups of organisms are unlikely to be genetically and demographically exchangeable (sensu (Templeton 1989)). Thus, sympatric and synchronous, discrete groups of organisms are widely accepted as relatively uncontroversial evidence of distinct, real units in nature (Mayr 1992; Coyne and Orr 2004; De Queiroz 2007; Mallet 2007, 2008).

The second issue is the degree to which taxonomic classification at the species level accurately reflects distinct groups of individual organisms or constitute arbitrary divisions of biological diversity (Rieseberg et al. 2006). To address this second issue about the nature of species, it is paramount to distinguish the idea of species taxa (i.e., taxonomic divisions at the species level) from the notion of species as biological units in nature (Hey et al. 2003). Species taxa do exist in the trivial sense of being human-made divisions, documented in taxonomic treatments. Yet, species may not occur in nature as discrete, sympatric and synchronous groups of individual organisms (Levin 1979; Raven 1986; Bachmann 1998; Barraclough 2019). Moreover, even if species do occur in nature as discrete sympatric and synchronous groups of individual organisms, human-made taxonomic classification at the species level may not reflect such groups. By example, the terms “splitter” and “lumper” were already used in Darwin’s time to (often pejoratively) describe taxonomists, including botanists, who divided biological diversity too finely and too broadly into species taxa, respectively (Endersby 2009).

These two issues about the nature of species may be empirically addressed in the context of multiple, potentially contrasting species definitions (*sensu* (De Queiroz 1999)). In particular, species could be real units in terms of different properties acquired during lineage divergence, including ability to interbreed, ecological divergence and reciprocal monophyly among others. Despite the variety of possible species definitions, there has been enduring interest in the reality of species in terms of two criteria: phenotypic and genome-wide distinctiveness (Fig. 1, (Dobzhansky 1951; Grant 1957; Mayr 1963; Ehrlich and Raven 1969; Levin 1979; Gould 2002; Coyne and Orr 2004; Rieseberg et al. 2006; Mallet 2013; Barraclough 2019)). Particular interest in these two species criteria seems to derive, at least in part, from the influential view of species as well-integrated phenotypic and genomic units, adapted to their physical and biotic environment (Dobzhansky 1951; Mayr 1963, see pages 518−540 in Gould 2002). According to this view, in sympatry and synchrony most species are phenotypically distinct lineages, characterized by distinct allele frequencies throughout the genome. This idea, according to which species can be characterized as concordant phenotypic and genomic groups, corresponds to the conventional species model (Barraclough 2019). Species that fit this model are sometimes referred to as “good” species (Fig. 1a, (Templeton 1989; Allmon 2016)).

**Figure 1.**
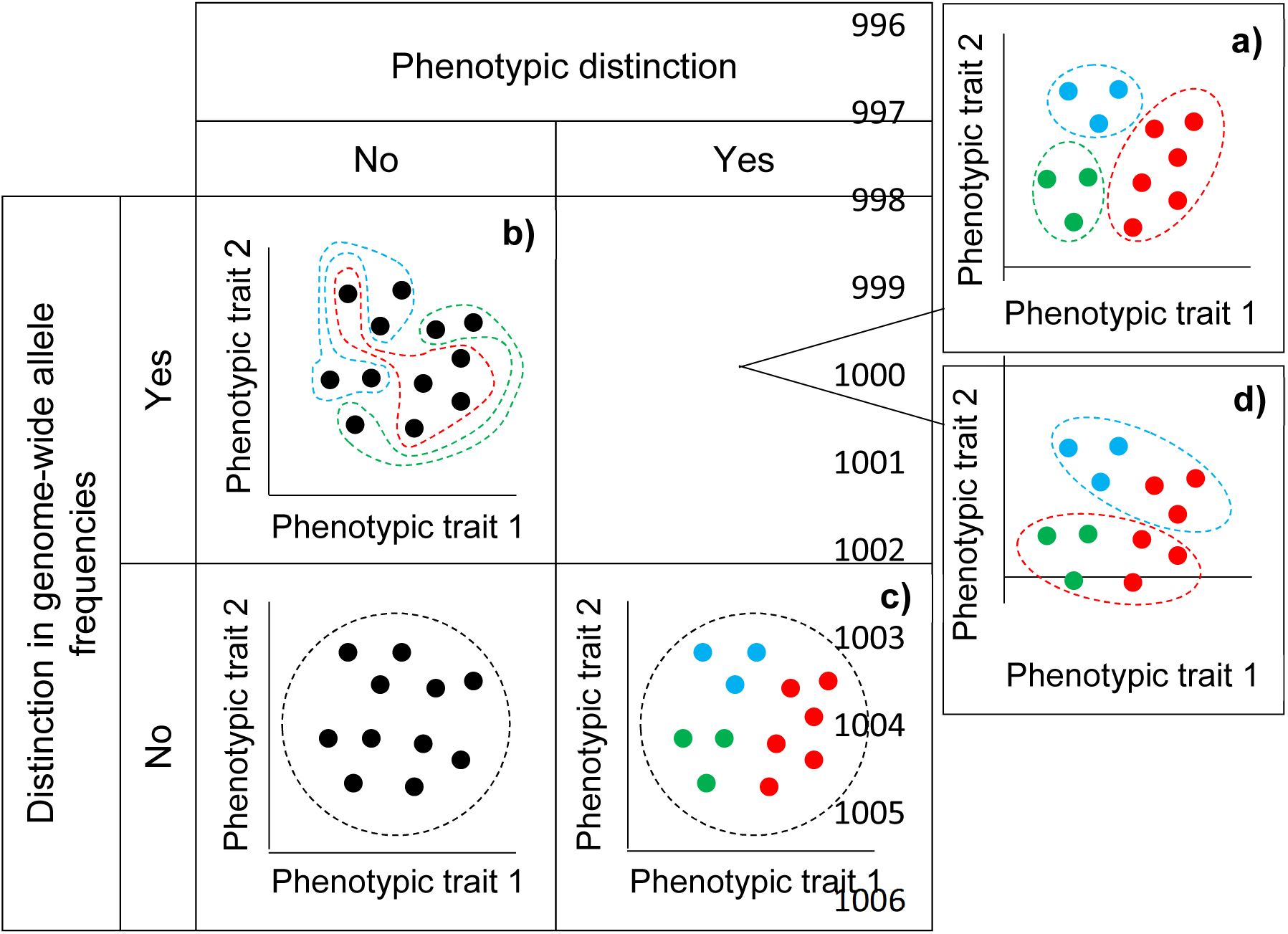
The nature of species in terms of phenotypic and genomic distinction. **a)** “Good” species are well-integrated units characterized by concordant phenotypic groups (colored points) and genomic groups (delimited by dashed lines). **b)** “Cryptic” species are phenotypically indistinct but differ in allele frequencies across many loci. **c)** Syngameons are characterized by distinct phenotypic groups in the absence of differences in allele frequencies across many or most loci. **d)** Distinct, discordant and non-nested phenotypic and genomic groups. In **b-d** phenotypes and genome-wide allele frequencies reflect contrasting evolutionary histories.

There are several alternatives to the conventional species model. Species may show very little if any phenotypic distinctiveness but differ in allele frequencies across many loci (Fig. 1b). These species are known as “sibling”(Mayr 1963) or “cryptic” species (Bickford et al. 2007; Fišer et al. 2018; Struck et al. 2018). Perhaps more controversial is the idea of species characterized by phenotypic distinctiveness in the absence of differences in allele frequencies across most loci (Fig. 1c), because it departs from the notion of species as well-integrated units (Wu 2001). Syngameons are examples of this latter kind of species, where phenotypic distinctiveness may be due to few loci, preserved despite interbreeding (Lotsy 1931; Grant 1971; Rieseberg and Burke 2001; Wu 2001) and the basis of meaningful ecological and evolutionary units (Van Valen 1976; Templeton 1989). Finally, some species may be characterized by discordant and non-nested sets of distinct phenotypes and genome-wide allele frequencies (Fig. 1d). In this latter case, as well as in cryptic species and syngameons, phenotypes and genome-wide allele frequencies might reflect contrasting evolutionary histories incorporated into a single lineage (Arnold 2015; Barraclough 2019).

While the degree of concordance between distinctiveness in phenotype and genome-wide allele frequencies (Fig. 1) have played a central role in debates about the nature of species, there seem to be few formal tests of whether coeval individual organisms form sympatric, distinct groups in nature according to these two criteria (Coyne and Orr 2004; Barraclough and Humphreys 2015; Barraclough 2019), and whether such groups (if any) correspond to species level taxonomic divisions (but see Rieseberg et al. 2006). Species delimitation studies would seem particularly well-poised to directly address these two aspects of the controversy about the nature of species. Several of these studies provide powerful insights by focusing on inference of genomic groups among individuals thought to represent different species taxa. Yet, because they are often not designed to measure phenotypes nor formally infer phenotypic groups, such studies do not test alternative species models in terms of concordance between distinctiveness in phenotype and genome-wide allele frequencies (Fig. 1). Particularly illustrative in this context are cases in which species are thought to be “cryptic” (Fig. 1b) because distinct genomic groups are found within species taxa. Given this kind of evidence, the claim that species are “cryptic” assumes concordance between species taxa (i.e., species level taxonomic divisions) and phenotypic groups. But such concordance is often poor (Rieseberg et al. 2006) and should ideally be tested as part of empirical studies on the nature of species (Fig. 2). In fact, morphological studies of presumed “cryptic” species often find that such species are actually phenotypically distinct (Allmon 2016; Korshunova et al. 2019).

**Figure 2.**
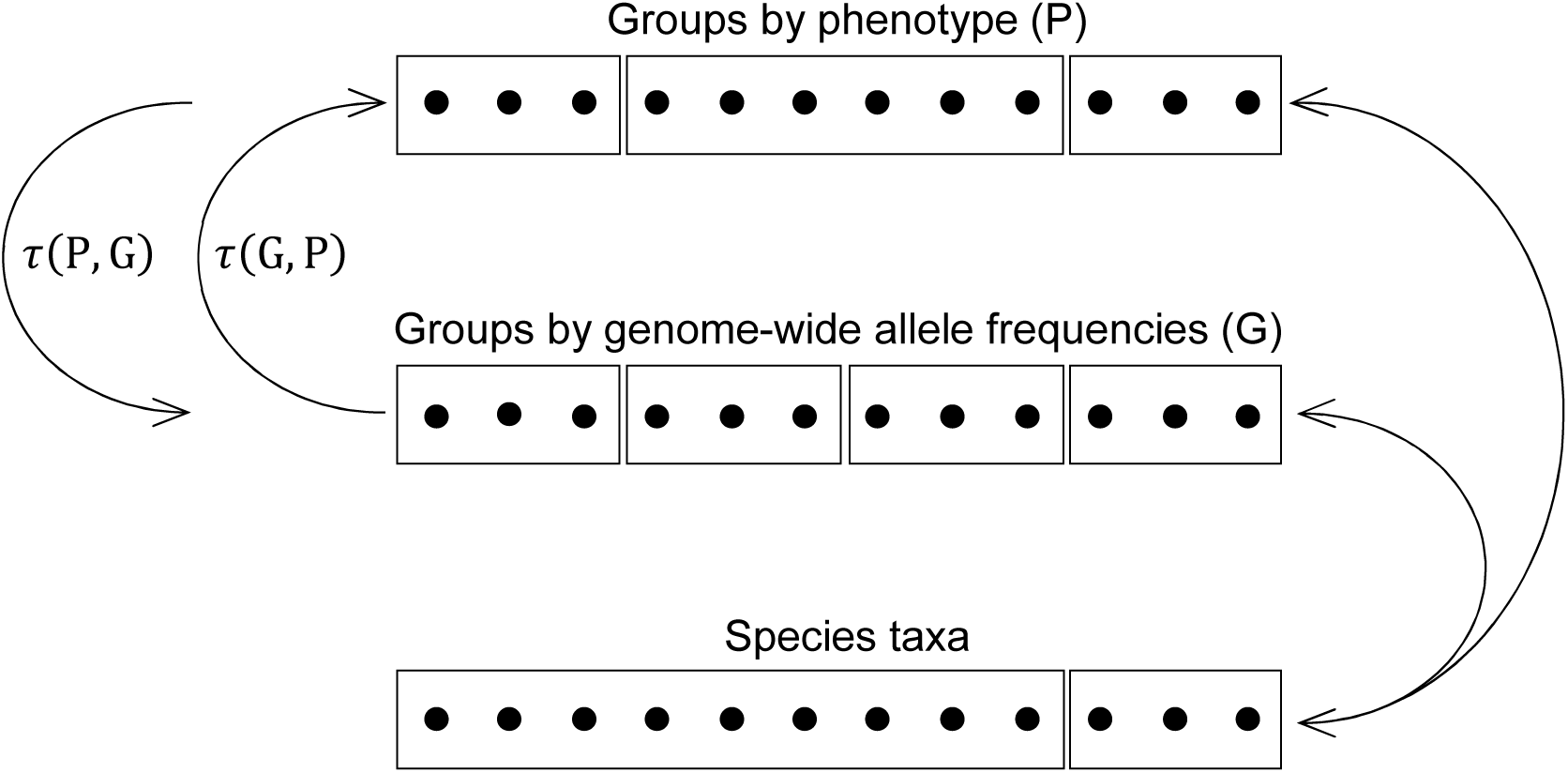
Illustrative example of degree of concordance between phenotypic groups, genomic groups and species taxa. Black dots represent a sample of sympatric and synchronic individuals, and rectangles represent groupings of this sample individuals according to different criteria. The top and middle rows of dots show phenotypic and genomic groups estimated *de novo*, respectively. The differences between phenotypic and genomic groups illustrates an instance of “cryptic” species (Fig. 1b). The lower row of dots illustrates an instance of species taxa defined by a “lumper.” The arrows on the left represent a single test of concordance between phenotypic and genomic groups. This test involves two values of the Goodman and Kruskal’s τ statistic, as illustrated by the two arrows on the left. In particular, τ(P,G) measures the ability to predict genomic groups based on information about phenotypic groups, while τ(G,P) measures the ability to predict phenotypic groups based on information about genomic groups (see equations 1 and 2). The double headed arrows on the right represent two tests of concordance, one between species taxa and phenotypic groups, and the other between species taxa and genomic groups. For simplicity, only one arrow is shown for each of these latter tests, even though each entails two values of the Goodman and Kruskal’s τ statistic.

The paucity of empirical studies directly addressing the nature of species in terms of concordance between distinctiveness in phenotype and genome-wide allele frequencies may also partly reflect the recency of methods to formally infer the number of distinct phenotypic groups in a sample of individuals. This inference should ideally be *de novo* (Barraclough 2019), as opposed to validation of previously proposed species hypothesis. Yet, commonly used approaches to the analysis of phenotypic data for species delimitation, such as discriminant function analysis (McLachlan 2004), are designed to test for differences between previously defined groups but not to infer the existence of such groups (Cadena et al. 2018). Another common approach to the analysis of phenotypic data relies on hierarchical clustering algorithms, even though phenotypic variation at and below the species level is unlikely to be hierarchical. Thus, these algorithms force data into hierarchical groups irrespective of whether such groups exist in nature(Crisp and Weston 1993; Queiroz and Good 1997). This problem is avoided by studies using ordination methods, but then inference of groups is often based on unwarranted reduction of dimensionality and subjective visual inspection of bivariate plots (Cadena et al. 2018). It is only with the relatively recent availability of tools to fit normal mixture models (e.g.,(Scrucca et al. 2016)) that these issues can be avoided (Ezard et al. 2010; Cadena et al. 2018). By contrast, *de novo* inference of genomic groups was widely adopted nearly two decades ago, soon after the development of ancestry models (Pritchard et al. 2000). These ancestry models, and subsequent refinements in their application (e.g.,(Verity and Nichols 2016, Lawson et al. 2018)), hold great potential for tests of species models defined in terms of concordance between phenotypic and genomic groups (Fig. 1). Yet, for that potential to be realized, approaches to *de novo* inference of phenotypic groups need to be widely adopted too.

Yet another reason for the scarcity of studies leveraging modern data and methods to directly address questions about the nature of species among synchronous and sympatric organisms may relate to the current emphasis in systematic biology on studies designed to sample the whole geographic distribution of a given monophyletic group. Due to virtually inexorable logistical limitations, such emphasis results in species delimitation studies that rely on geographically extensive and sparse sampling of organisms, as opposed to more thorough samples of the organism co-occurring within localities or regions. The relative merits of these two approaches to sampling, in terms of insights into species delimitation, was a central issue for salient botanists of the XIX century that closely interacted with Darwin, including Asa Gray and Joseph Dalton Hooker (Stevens 1997). In practice the issue seems to have been resolved siding with Hooker, who emphasized geographically extensive samples of closely related groups of organisms rather than intensive regional samples. Thus, the sampling strategy of modern botanical collectors may be characterized as “never the same species twice” (ter Steege et al. 2011), in reference to their (intended) behavior in a single collecting locality. Currently, this approach to sampling is widely practiced among non-botanists too, and results in a characteristic pattern in the locality data derived from natural history collections: well-separated collection localities and few specimens per locality (Sheth et al. 2012). No doubt, geographically extensive samples have contributed dramatically to our understanding of species limits. Nevertheless, geographically sparse sampling is insufficient to test species models, as defined in terms of phenotypic and genomic groups (Fig. 1), with the relatively uncontroversial evidence provided by studies of synchronic and sympatric organisms.

In sum, despite noteworthy efforts (e.g., (Cavender-Bares and Pahlich 2009)), the debate about the nature of species remains unsatisfactorily addressed by empirical studies, due at least in part to the scarcity of germane analysis of phenotypic data and the emphasis of species delimitation studies on geographically extensive rather than locally intensive sampling of individual organisms. Here we contribute to the empirical resolution of the debate by testing alternative species models (Fig. 1) and, also, by assessing the degree to which taxonomic divisions at the species level (i.e., species taxa) accurately reflect the phenotypic and genomic distributions of coeval, sympatric individual organisms (Fig. 2). We aim to present a case study characterized by geographically dense sampling of organisms, and the use of formal approaches to i) *de novo* inference of groups of organisms in terms of phenotype and genome-wide allele frequencies, ii) measure the degree of concordance between distinctiveness in phenotype and genome-wide allele frequencies, and iii) the concordance between species taxa and distinctiveness in phenotype and genome-wide allele frequencies. We are unaware of empirical studies about the nature of species that employ these approaches on a geographically dense sampling of individual organisms. Yet, such studies are needed to empirically settle the debate about the nature of species.

We focused on *Espeletia*, a genus in the plant subtribe Espeletiinae (Asteraceae) that is endemic to the northern Andes (Monasterio and Sarmiento 1991; Rauscher 2002; Cuatrecasas 2013; Diazgranados and Barber 2017) and often dominant in high elevation ecosystems known as “páramos” (Ramsay and Oxley 1997). This genus appears to have undergone a very rapid radiation starting around 2.7 Ma (Madriñán et al. 2013; Pouchon et al. 2018). *Espeletia* species are suspected to commonly produce inter-specific hybrids, even between distantly related species (Pouchon et al. 2018), and, nevertheless, maintain phenotypic and ecological integrity (Berry et al. 1988; Diazgranados 2012; Cuatrecasas 2013; Diazgranados and Barber 2017). In one of the (two) prefaces of the most recent Espeletiinae monograph (Cuatrecasas 2013), Harold Robinson wrote: “The recent origin, the high specialization and the complex structure of the often large plants, added to the few barriers to hybridization in addition to geography, even at the intergeneric level, presented many complications for the monographic study.” Indeed, along with other plant genera such as oaks (*Quercus*), hawthorns (*Crategus*) and blackberries (*Rubus*), *Espeletia* may be a “botanical horror story” (Diamond 1992) from which we may learn much about the nature of species.

## Materials AND Methods

### Study Region and Taxa

We studied plants of the genus *Espeletia* occurring in the páramo de Sumapaz. This páramo encompasses 1,780 km^2^ in the Cordillera Oriental of the Colombian Andes, south of Bogotá, and includes the high elevation areas of Parque Nacional Sumapaz. Annual average temperature varies spatially between 2 and 10 °C and annual rainfall between 500 and 2000 mm. According to the most recent *Espeletia* monograph (Cuatrecasas 2013), seven species taxa in the genus *Espeletia* occur in the páramo de Sumapaz: *Espeletia argentea* Humb. & Bonpl., *Espeletia cabrerensis* Cuatrec., *Espeletia grandiflora* H. & B., *Espeletia killipii* Cuatrec., *Espeletia miradorensis* (Cuatrec.) Cuatrec., *Espeletia summapacis* Cuatrec. and *Espeletia tapirophila* Cuatrec. The monograph also explicitly mentions the occurrence of infra-specific taxa in the páramo de Sumapaz for three of the seven species taxa: *Espeletia argentea fma. phaneractis* (S.F.Blake) A.C.Sm., *E. grandiflora spp. grandiflora var. attenuata* Cuatrec., *E. grandiflora spp. subnivalis* Cuatrec. and *Espeletia killipii var. chiscana* Cuatrec.

### Specimen Collection

To sample *Espeletia* species we defined the limit of the páramo de Sumapaz as the 3,000 m elevation isocline, based on the Aster digital elevation model with 30 m resolution (https://asterweb.jpl.nasa.gov/gdem.asp). Thus defined, the páramo de Sumapaz was divided into quadrants of 2 x 2 km. We sampled 34 of these quadrants by randomly choosing two 30 × 30 m grid cells (of the Aster digital elevation model) in each band of 100 m elevation (Fig 3). We sampled a total of 219 grid cells of 30 × 30 m. In each grid cell we collected at least one specimen of each seemingly different phenotype. Additionally, we collected specimens opportunistically to further document the occurrence of different phenotypes across the páramo de Sumapaz. In total we collected 538 specimens of reproductive *Espeletia* plants, all deposited in triplicate at the herbarium of the Jardín Botánico de Bogotá José Celestino Mutis (JBB).

**Figure 3.**
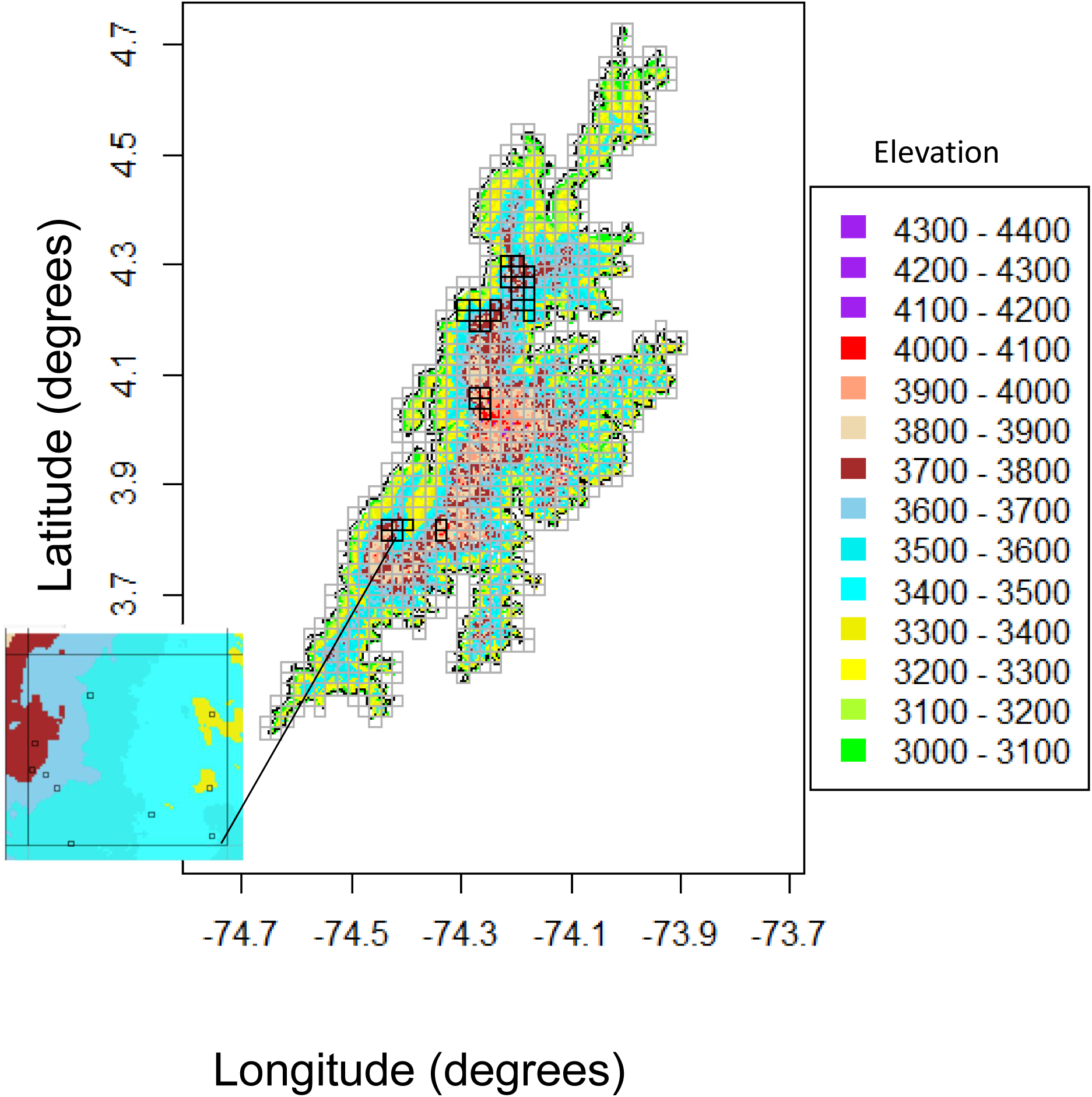
Topographic map of the study region, delimited by the contour of 3,000 m of altitude, according to the digital Aster elevation model (https://asterweb.jpl.nasa.gov/gdem.asp). The study region was divided into quadrants of 2 × 2 km (gray grid). We sampled 34 of these quadrats (black squares). In particular, in each of these 34 quadrants we sampled two 30 × 30 m cells in each band of 100 m altitude, chosen randomly based on the Aster digital elevation. For example, the 2 × 2 km quadrant shown to the left of the figure has five elevation bands (as indicated by the color scale). Therefore, we sampled 10 cells of 30 × 30 m, two in each elevation band of 100 m, as indicated by the black squares within the 2 × 2 km quadrant.

### Phenotypic Distinctiveness

### Morphological characters

We measured thirteen morphological characters in *Espeletia* specimens from the páramo de Sumapaz (Table 1). Except perimeter of the synflorescence axis, all these characters were used by Cuatrecasas (2013) to delimit *Espeletia* species taxa from the páramo de Sumapaz. We additionally included the perimeter of the synflorescence axis because in the field this character seemed useful to distinguish putative species. Whenever possible, we measured all thirteen characters in each of the 538 specimens we collected. Additionally, we measured as many of the thirteen characters as possible in each of 165 specimens from páramo de Sumapaz, or from taxa known to occur there, that were collected by other researchers and deposited in the Herbario Nacional Colombiano (COL), at the Instituto de Ciencias Naturales, Universidad Nacional de Colombia. Out of these 165 specimens, 97 were explicitly mentioned in the *Espeletia* monograph by Cuatrecasas (2013), including types for all but one (*Espeletia argentea*) species taxa that, according to Cuatrecasas (2013), occur in the páramo de Sumapaz.

**Table 1.** Thirteen morphological characters measured in *Espeletia* specimens from the páramo de Sumapaz to test for the existence of phenotypic groups.

| Character number and name | Character description |
| --- | --- |
| 1) Leaf blade length | Distance from the apex to the base of the leaf blade (excluding the leaf sheath). |
| 2) Leaf blade width | Width of the leaf blade at the widest part. |
| 3) Number of capitula per synflorescence | Count of the number of capitula in a single synflorescence. |
| 4) Capitulum diameter | Diameter of the capitulum, omitting the ligules of the flowers of the radius. |
| 5) Number of pairs of sterile leaves per synflorescence | Count of the pairs of leaves in the infertile (or vegetative) part of the synflorescence axis. |
| 6) Length of the synflorescence axis | Distance from the base of the central capitulum in the terminal cyme to the base of the synflorescence axis. |
| 7) Synflorescence axis perimeter | Perimeter around the synflorescence axis at 10 cm from the base (avoiding areas near sterile leaves). |
| 8) Terminal cyme peduncle length | Length of the peduncle of the central capitulum in the terminal cyme. |
| 9) Sterile phyllary length | Distance from the apex to the base of the sterile phyllary subtending capitula. |
| 10) Sterile phyllary width | Width of the sterile phyllary subtending capitula, at the widest part. |
| 11) Length of the corolla of the flowers in the disc | Distance from the distal tip of a corolla lobe to the base of the tube of the flowers in the disc. |
| 12) Length of the corolla of the flowers in the ray | Distance from the distal tip of the ligule to the base of the tube of the flowers in the ray. |
| 13) Length of the tubes of the flowers in the disc | Distance from the base of the tube to the widening of the corolla of the flowers in the disc. |

When possible, each one of the thirteen morphological characters was measured in three separate structures per specimen (as mentioned before, specimens were collected in triplicates), and analyses were based on average values per specimen to account for intra-individual variation. The length of synflorescence axis was measured in the field to the nearest centimeter. All other measurements were done in the herbarium after the specimens were dried. To measure leaf blade length and width we photographed leaves on a standard background including a millimeter ruler and then processed the respective images using *Image J* (Schindelin et al. 2012). Capitulum diameter, length and width of the corolla of the flowers of the disc, length of the corolla of the flowers of the ray, and length of the tubes of the disc were measured to the nearest hundredth of a millimeter. All morphological data is available as Supplementary Material in Appendix 1.

### Estimating morphological groups

To estimate *de novo* the number phenotypic groups among the measured *Espeletia* specimens, as well as the assignment of specimens to such groups, we used the R package Mclust 5.0 (Scrucca et al. 2016) to fit normal mixture models (NMMs, (McLachlan and Peel 2000)) to morphological data. The use of NMMs to detect phenotypically distinct species assumes polygenic inheritance of phenotypic traits and random mating; under these assumptions, gene frequencies would be close to Hardy-Weinberg equilibrium, two or more loci would be near linkage equilibrium, and phenotypic variation among individuals of a single species would tend to be normally distributed (Templeton 2006). Thus, detection of two or more phenotypic normal distributions among sympatric and synchronic organisms indicates the existence of two or more phenotypically distinct species (Cadena et al. 2018), barring instances in which distinct normal distributions reflect ontogenetic variation or phenotypic plasticity. The parameters of NMMs (e.g., means, variances and covariances) describe each phenotypic group as a (possibly multivariate) normal distribution and can be estimated *de novo* from data on phenotypic measurements (i.e., without *a priori* knowledge of species limits) using the expectation-maximization algorithm (McLachlan and Krishnan 2008).

We used the Bayesian Information Criterion (BIC, (Schwartz 1978)) to measure empirical support for different NMMs fitted to the morphological data (Fraley and Raftery 2002). We fitted NMMs that covered a wide range of the possible number of distinct phenotypic groups, from a single group up to ten groups. This range amply bracketed the number species taxa that, according to the most recent monograph of *Espeletia* (Cuatrecasas 2013), occur in the páramo de Sumapaz: seven (see above section on *Study Region and Taxa*). Before fitting MMNs, we rotated the phenotypic space using a principal component analysis on the covariance matrix of the natural logarithm of the measurements of thirteen morphological characters (Table 1). One of these characters, pairs of sterile leaves per synflorescence, took on zero values, so we added one to this variable before the logarithm transformation. We used R package Clustvarsel (Scrucca and Raftery 2018) to reduce the dimensionality of the data by selecting the most useful set of principal components for the discrimination of groups in NMMs, without *a priori* knowledge of the groups (Raftery and Dean 2006; Maugis et al. 2009). We separately used algorithms implemented in Clustvarsel that performed backward and forward selection of principal components.

To avoid convergence of the expectation maximization algorithm on a local maximum of the likelihood function, we ran five versions of the analyses based on NMMs, including reduction of dimensionality with package Clustvarsel, each using a different approach for the initialization of the expectation maximization algorithm (Scrucca and Raftery 2015). All five approaches to initialization are available in the package Mclust. Each approach applies a different data standardization to obtain an initial partition of the data via hierarchical agglomerative clustering (Scrucca and Raftery 2015), although the reminder of the expectation maximization algorithm is run on the data on the initial scale (i.e., the principal components of the covariance matrix of the natural logarithm of the morphological measurements). Details of the analysis based on NMMs implemented in package Mclust, including reduction of dimensionality using package Clustvarsel, are in the R scripts available as Supplementary Material in Appendix 2.

### Distinctiveness in Genome-wide Allele Frequencies

#### DNA extraction and genotyping-by-sequencing

Genomic DNA was extracted using the QIAGEN DNeasy Plant Mini Kit (QIAGEN, Germany). DNA concentration was quantified in R Qubit dsDNA HS Fluorometer (Life Technologies, Sweden). One 96-plex genotyping-by-sequencing libraries were performed with MsII digestions (Elshire et al. 2011) and sequenced at LGC Genomics (Berlin, Germany). Readings were aligned based on a previous study (Cortés et al. 2018) using a cluster of 283 *Espeletia* specimens, built with 192 specimens from that study in addition to 96 specimens from this study. Filtering of variants was done using a GBS-specific rule set with > 5 read count for a locus, > 5% minimum allele frequency across all specimens, and ≥ 80% genotypes observed. This filtering reduced the initial sample of 99 specimens to 77. The filtered dataset was inspected with TASSEL v. 3.0 (Glaubitz et al. 2014), resulting in a final set of 2,098 single nucleotide polymorphisms (SNPs).

### Estimating groups according to genome-wide allele frequencies

To estimate *de novo* the number of groups according to genome-wide allele frequencies among genotyped *Espeletia* specimens, and assign specimens to such groups, we implemented ancestry models in *R Maverick* package in R 3.3.1. (Verity and Nichols 2016). These models assume that groups tend to be in the Hardy Weinberg equilibrium with their own set of allele frequencies and that all loci are in linkage equilibrium (Pritchard et al. 2000). Yet, all ancestry models we used allowed admixture. We evaluated ancestry models postulating a wide range of number of groups among *Espeletia* specimens, from a single group up to ten. Again, this range includes the number of species taxa that, according to the most recent monograph of *Espeletia* (Cuatrecasas 2013), occur in the páramo de Sumapaz. R Maverick assesses the number of groups using thermodynamic integration to obtain direct estimates of the posterior probability of models, given the data and a prior distribution of model parameters (Verity and Nichols 2016).

### Species Taxa

To determine the extent to which species taxa corresponded to phenotypic groups and groups based on genome-wide allele frequencies, we assigned specimens to species taxa based on detailed descriptions in Cuatracasas (2013). In particular, for all species taxa expected to occur in Sumapaz according to Cuatrecasas (2013), we obtained the maximum and minimum values for twelve of the thirteen morphological characters in Table 1. We excluded from this analysis one of the measured characters, perimeter of the synflorescence axis, because it was not considered by Cuatrecasas (2013), as already mentioned. Then, for each specimen in our sample, we determined if the measured values for the twelve characters fell within the ranges of each species taxa. We assigned a specimen to a species taxon if the measurements for all twelve characters fell within the respective ranges provided by Cuatracasas (2013).

### Concordance Between Groups Defined by Phenotype, Genome-wide Allele Frequencies and Species Taxa

To estimate the degree of concordance between the groups of specimens defined according to phenotypes, genome-wide allele frequencies and species taxa we used Goodman and Kruskal’s tau statistic (τ, (Agresti 2002)) as implemented in R package GoodmanKruskal version 0.0.2 (Pearson 2016). This statistic is based on measuring variability as the probability that two items (specimens in our case) selected at random belong to different groups. Thus, for an assignment of specimens to groups according to phenotypes (P), variability equals the probability that two randomly drawn specimens belong to different phenotypic groups, hereafter V(P). Likewise, for an assignment of specimens to groups according to genome-wide allele frequencies (G), variability is the probability that two randomly drawn specimens belong to different genomic groups, hereafter V(G).

In contrast to other metrics of association between categorical variables (e.g., chi-square, Cramer’s V), τ is a potentially asymmetric measure of association (Agresti 2002). Thus, τ(P,G) measures the ability to predict genomic groups based on information about phenotypic groups, while τ(G,P) measures the ability to predict phenotypic groups based on information about genomic groups (Fig. 2):

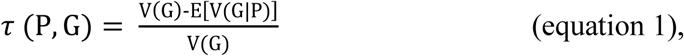

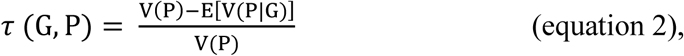

where V(G|P) is the probability that two specimens randomly drawn from a single phenotypic group belong to two different genomic groups, and V(P|G) is the probability that two specimens randomly drawn from a single genomic group belong to two different phenotypic groups. E[V(G|P)] and E[V(P|G)] are expected values of these quantities across phenotypic and genomic groups, respectively. The values of τ(P,G) and τ(G,P) range between zero and one. Values near one (zero) indicate high (low) predictive ability.

Asymmetry between τ(P,G) and τ(G,P) may arise because phenotypic groups may predict genomic groups better than genomic groups may predict phenotypic groups, or vice versa. Crucially, the different species models in Figure 1 imply particular asymmetries, or absence of asymmetry between τ(P,G) and τ(G,P). “Good” species imply strong concordance between phenotypic and genomic groups (Fig. 1a) and, therefore, little if any asymmetry between τ(P,G) and τ(G,P). Both values should be high and statistically significant. In other words, phenotypic groups should predict genomic groups well and vice versa. In contrast, among “cryptic” species genomic groups should predict phenotypic groups better than phenotypic groups can predict genomic groups, given that genomic groups are nested within phenotypic groups (Fig. 1b). Thus, for “cryptic” species τ(P,G) < τ(G,P). The opposite is expected in syngameons because phenotypic groups are nested within genomic groups (Fig. 1c) and, therefore, phenotypic groups should predict genomic groups better than phenotypic groups can predict genomic groups. Finally, in the case of discordant and non-nested sets of distinct phenotypes and genome-wide allele frequencies (Fig. 1d), values of τ(P,G) and τ(G,P) would both be low and lack statistical significance.

Similar expectations can be described for concordance, as measured by τ, between species taxa and phenotypic or genomic groups (Fig. 2). By example, when species taxa correspond to phenotypic or genomic groups, τ values would be high, statistically significant and symmetric. On the other hand, species taxa defined by “lumpers” would each include several phenotypic or genomic groups. Given this nesting, the ability to predict phenotypic or genomic groups based on species taxa would be low, even though the ability to predict species taxa from information on phenotypic or genomic groups would be high. The opposite would be expected for species taxa defined by “splitters.” Finally, in the case of non-nested discordance between species taxa and phenotypic or genomic groups, all τ values would be low and lack statistical significance.

We evaluated the statistical significance of τ values using 1,000 iterations of a null model that randomized the assignment of specimens to groups in one of the two categorical variables being compared (phenotypic groups, genomic groups or species taxa, Fig. 2). We calculated τ values for each iteration of the null model and thus constructed a null distribution of 1,000 values. We estimated p-values for each test as the fraction of τ values in the null distribution that were at least as extreme as the observed τ value.

### Sympatry

To assess whether phenotypic and genomic groups were sympatric, we adopted an operational definition of sympatry reflecting both potential for gene flow and geography (Mallet et al. 2009). According to this definition, populations are sympatric when they have geographic distributions within the normal cruising range of individuals of each other (Mayr 1947), so that gametes of individuals are physically capable of encountering one another with moderately high frequency (Futuyma and Mayer 1980). The normal cruising range of individual organisms is determined by the distribution of distances between the sites of birth and breeding (Mallet et al. 2009). Thus, both pollen and seed dispersal are relevant to this operational definition of sympatry, although in *Espeletia* seed dispersal is thought to be fairly limited (Berry and Calvo 1989; Diazgranados 2012; Gallego Maya and Bonilla Gómez 2016). In sharp contrast, the pollen of *Espeletia* can travel considerable distances via bumblebees (genus *Bombus*) and hummingbirds (Fagua and Gonzalez 2007); except in some species that seem to be pollinated by wind (Berry and Calvo 1989). Bumblebees home back to their nests from places 9.8 km away (Goulson and Stout 2001) and regularly perform flights covering > 1 km (Greenleaf et al. 2007; Pope and Jha 2018). Thus, we considered inter-group distances < 1 km as small enough for sympatry. Accordingly, to estimate the degree of sympatry between a given pair of groups (phenotypic or genomic), we measured the geographic distance between each specimen and the nearest specimen that did not belong to the same group. We examined the resulting distribution of inter-group distances to determine if groups were separated by < 1 km.

## RESULTS

### Phenotypic Distinctiveness

The morphological analysis included only 307 out of 703 specimens measured in this study, because several specimens were missing values for at least one of the 13 characters studied (Table 1), most often flower traits. The procedure for dimensionality reduction (Clustvarsel, see Methods) selected the first 12 principal components, out of a total of 13, for discrimination of morphological groups in normal mixture models (NMMs). This result was obtained regardless of whether the algorithm for reduction of dimensionality employed a forward or backward search of principal components.

We found distinct phenotypic groups among the *Espeletia* specimens from páramo de Sumapaz. In particular, the best NMM identified six morphological groups and had substantially more empirical support than models assuming 1−5 and 7−10 groups (ΔBIC > 10, Fig. 4a). In this best model, the morphological groups appear to be fairly distinct, because assignment of specimens to morphological groups entailed little uncertainty. In a probability scale, assignment uncertainty exceeded 0.1 in only 3 out of 307 specimens.

**Figure 4.**
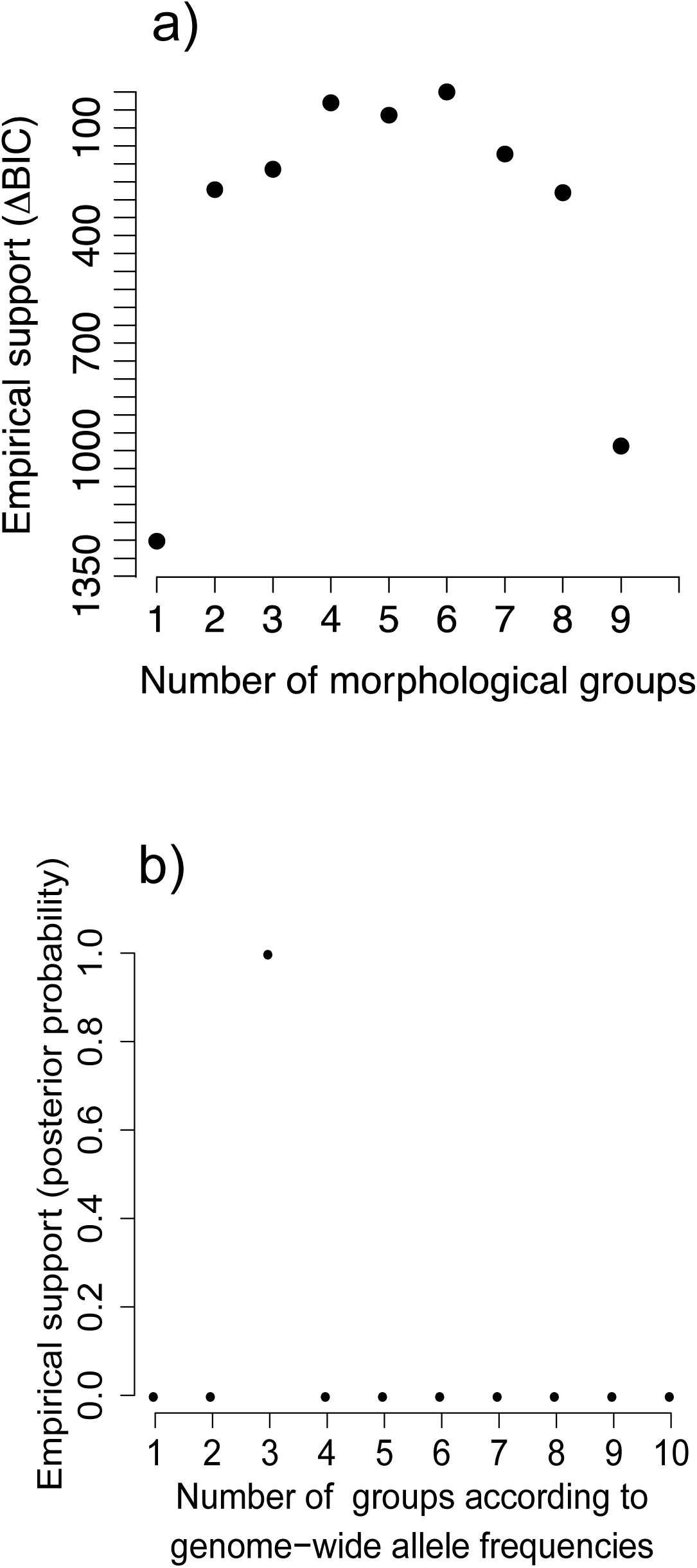
Empirical support for **a)** normal mixture models assuming different number of morphological groups and **b)** ancestry models assuming different number of groups defined by genome-wide allele frequencies among *Espeletia* from páramo de Sumapaz. The normal mixture model with highest empirical support has 6 morphological groups while the best ancestry model distinguishes 3 genomic groups.

In the best NMM model, three groups were relatively easily distinguished in the morphological space defined by the first two principal components, mainly determined by four characters: number of capitula per synflorescence, number of sterile leaves per synflorescence, length of the cyma peduncle and sterile phyllary width (Fig. 5a). At one extreme of this space, morphological group 1 was distinguished from the rest by having few capitula per synflorescence and no sterile leaves along the synflorescence (extreme right in Fig. 5c). At another extreme, morphological group 4 had the narrowest sterile phyllaries, numerous capitula per synflorescense and the shortest cyma peduncles (upper left in Fig. 5c). Finally, morphological group 6 can be distinguished from the rest by wide sterile phyllaries, many capitula and several pairs of sterile leaves per synflorescence (central lower part in Fig. 5c). The remaining morphological groups were better distinguished in the morphological space defined by principal components 2 and 3, mainly determined by leaf blade width, number of capitula per synflorescence, number of sterile leaves per synflorescence, sterile phyllary width and length of the tubes of the flowers in the disc (Fig. 5b). In this space, morphological groups 2, 3 and 5 were located along a gradient of decreasing values for the number of sterile leaves per synflorescence (Fig. 5d).

**Figure 5.**
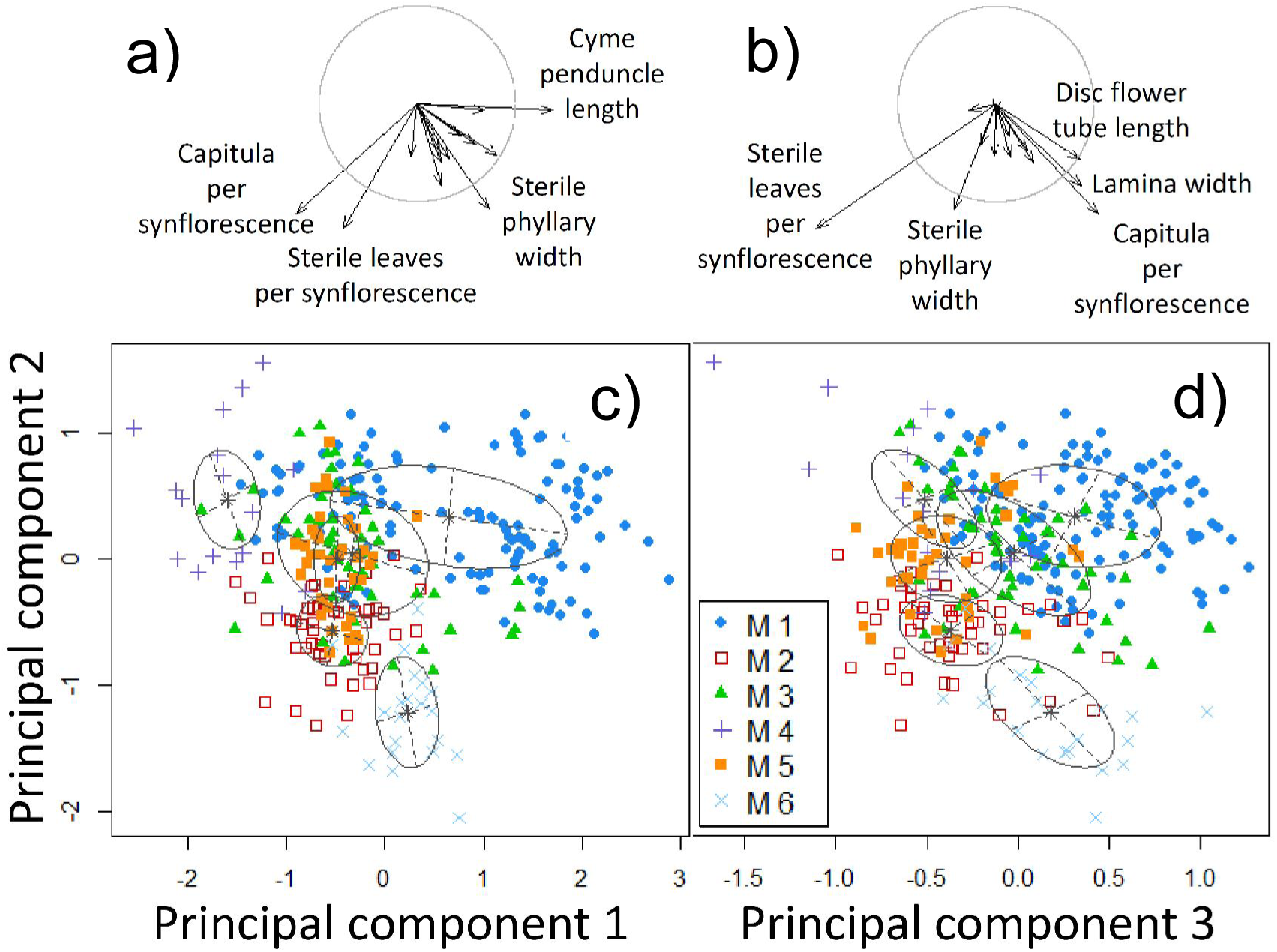
Six phenotypic groups detected by normal mixture models among *Espeletia* specimens from paramo de Sumapaz, as seen in the morphological space dtermined by the first 3 principal components. A subset of 6 characters largely determined these 3 principal components (**a, b**), where phenotypic groups can be distinguished (**c, d**). Arrows in **a** and **b** display the contribution of characters to principal components (loadings), and circles show the expected length of arrows if all characters contributed equally. Arrows exceeding this expectation contribute most significantly and are labeled. In **c** and **d**, ellipses show regions within 1 standard deviation of the multivariate mean of each phenotypic group, and symbols represent specimens according to morphological groups (M1 to M6).

### Distinctiveness in Genome-Wide Allele Frequencies

We found support for the existence of more than one genomic group among the *Espeletia* plants we sampled from the paramo de Sumapaz. In particular, an ancestry model with three groups had the highest empirical support, with posterior probability of almost one (Fig. 4b). In this best ancestry model, however, the genomic groups did not seem particularly distinct. One of these groups had only six specimens, all with a relatively low proportion of alleles from other groups (Fig. 6a). The other two groups were much larger (32 and 39 specimens) and included specimens with fairly high proportions of mixed ancestry. Indeed, in both these groups specimens with more than 10% admixture were nearly half the group (19 and 21, respectively), and fairly high values of admixture (> 20%) were not uncommon (Fig. 6a). Moreover, six specimens with admixture values near 50% suggest F1 hybrids among the three groups are not uncommon.

**Figure 6.**
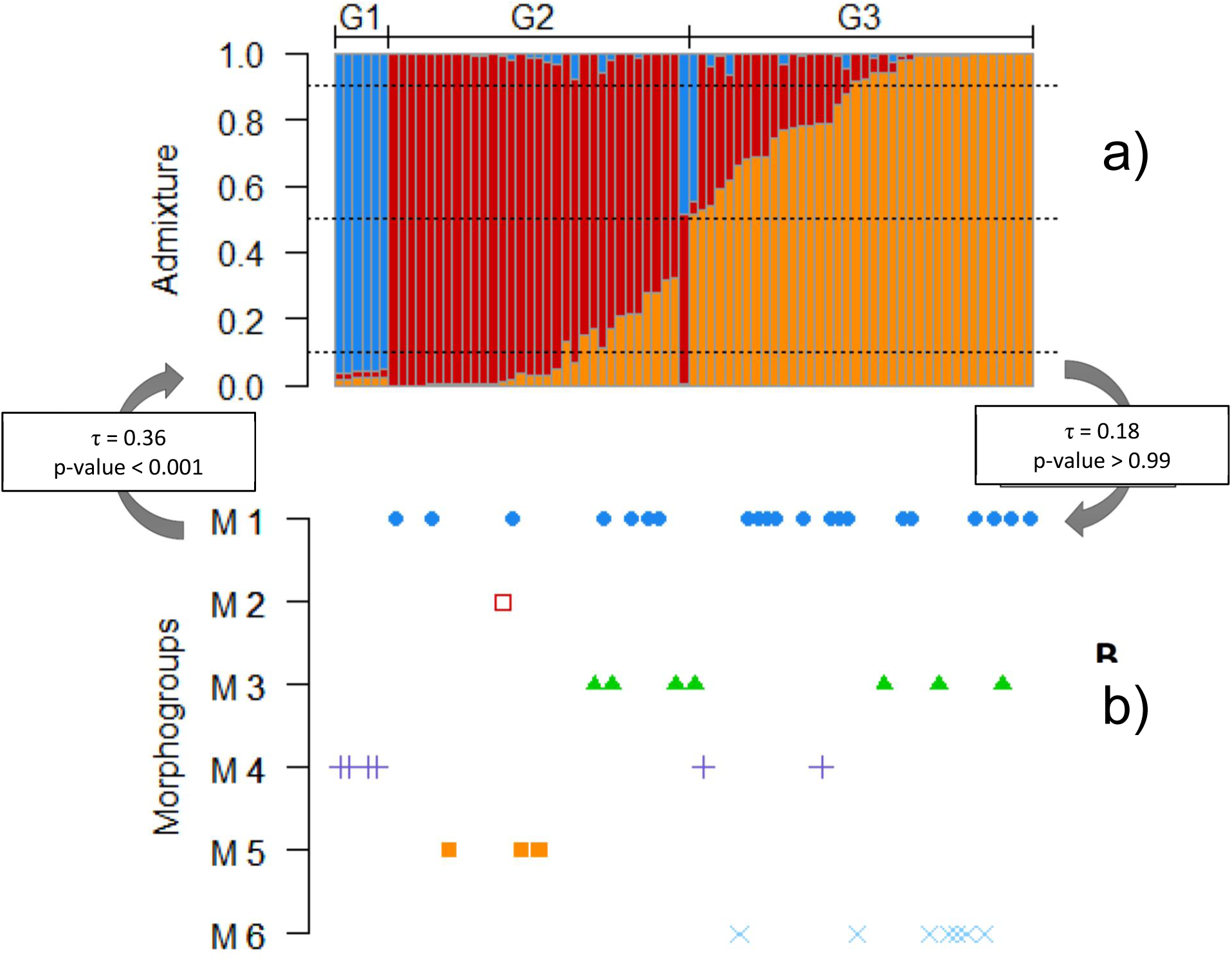
Three genomic groups detected by ancestry models among *Espeletia* specimens from paramo de Sumapaz and concordance with phenotypic groups detected by normal mixture models (Fig. 5). **a)** assignment of specimens to genomic groups. Each specimen is represented by a stacked bar showing admixture proportions in color. Labels at the top indicate specimen assignment to genomic groups (G1 to G3). **b)** Assignment of specimens to morphological groups (vertical axis, symbols as in Fig. 5). Some specimens in the gnomic groups could not be classified into phenotypic groups because they were missing morphological characters. There was low concordance between genomic and morphological groups, as seen in low values of the Goodman and Kruskal tau statistic (τ).

### Species Taxa

The procedure to assign specimens to species taxa described by Cuatrecasas (2013) resulted in only one specimen being assigned to any species taxa, out of a total of 307 specimens with data for all 12 relevant morphological characters (i.e., all characters in Table 1 except perimeter of the synflorescence axis). In particular, one specimen collected during field work for this study was assigned to *Espeletia killipii* Cuatrec. All other specimens fell outside the ranges specified by Cuatrecasas (2013) for the species expected to occur in the páramo de Sumapaz, including 27 specimens explicitly mentioned by Cuatrecasas (2013). Among them are specimens labeled (i.e., determined) as *E. argentea, E. grandiflora, E. killipii* and *E. summapacis* and the type of *E. tapirophila*.

### Concordance Between Groups Defined by Phenotype, Genome-wide Allele Frequencies and Species Taxa

The remarkable result presented in the previous section, whereby all but one of the 307 *Espeletia* specimens in our sample could not be formally assigned to any species taxa, meant that we could not use the Goodman and Kruskal’s tau statistic (τ, see Methods) to estimate the degree of concordance between species taxa and groups defined by phenotypic distinctiveness or genome-wide allele frequencies. This statistic assumes that specimens are actually assigned to groups, but the morphological space defining species taxa (according to Cuatrecasas 2013) was nearly void of specimens. Nonetheless, it seems fair to state that concordance between species taxa and groups defined by phenotypic distinctiveness or genome-wide allele frequencies was null, because species taxa could not predict assignment of specimens to either phenotypic distinctiveness or genome-wide allele frequencies, and visce versa.

The analysis of concordance between the classifications based on phenotypic distinctiveness and genome-wide allele frequencies included 46 specimens, because several specimens in the genomic classification were not in anthesis and, therefore, could not be included in the phenotypic classification based on all morphological traits (Table 1). The association between these classifications was not very strong, although it was statistically significant in one direction. In particular, knowledge of phenotypic groups conferred statistically significant but poor ability to predict groups based on genome-wide allele frequencies (τ = 0.36, p-value < 0.001). This weak concordance (τ ranges from zero to one, see Methods) seems mainly due to the complete inclusion of morphological groups 2, 5 and 6 into single genomic groups (Fig. 6). However, each of the three remaining morphological groups straddled two genomic groups. The association between classifications was even weaker and not statistically significant when measured in terms of the ability to predict assignment of specimens to phenotypic groups based on knowledge about genomic groups (τ = 0.177, p-value > 0.001). This result reflects the inclusion of multiple phenotypic groups in each of the two genomic groups with most specimens (Fig. 6). We note, however, that the genomic group with the smallest number of specimens included only morphological group 4 (Fig. 6).

### Sympatry

All phenotypic and genomic groups were sympatric, given the assumption that inter-group geographic distances ≤ 1 km allowed gamete exchange via pollen dispersal by bumblebees. All geographic distances between pairs of morphological groups were ≤ 1 km, and the same was true for groups defined by genome-wide allele frequencies (Fig. 7).

**Figure 7.**
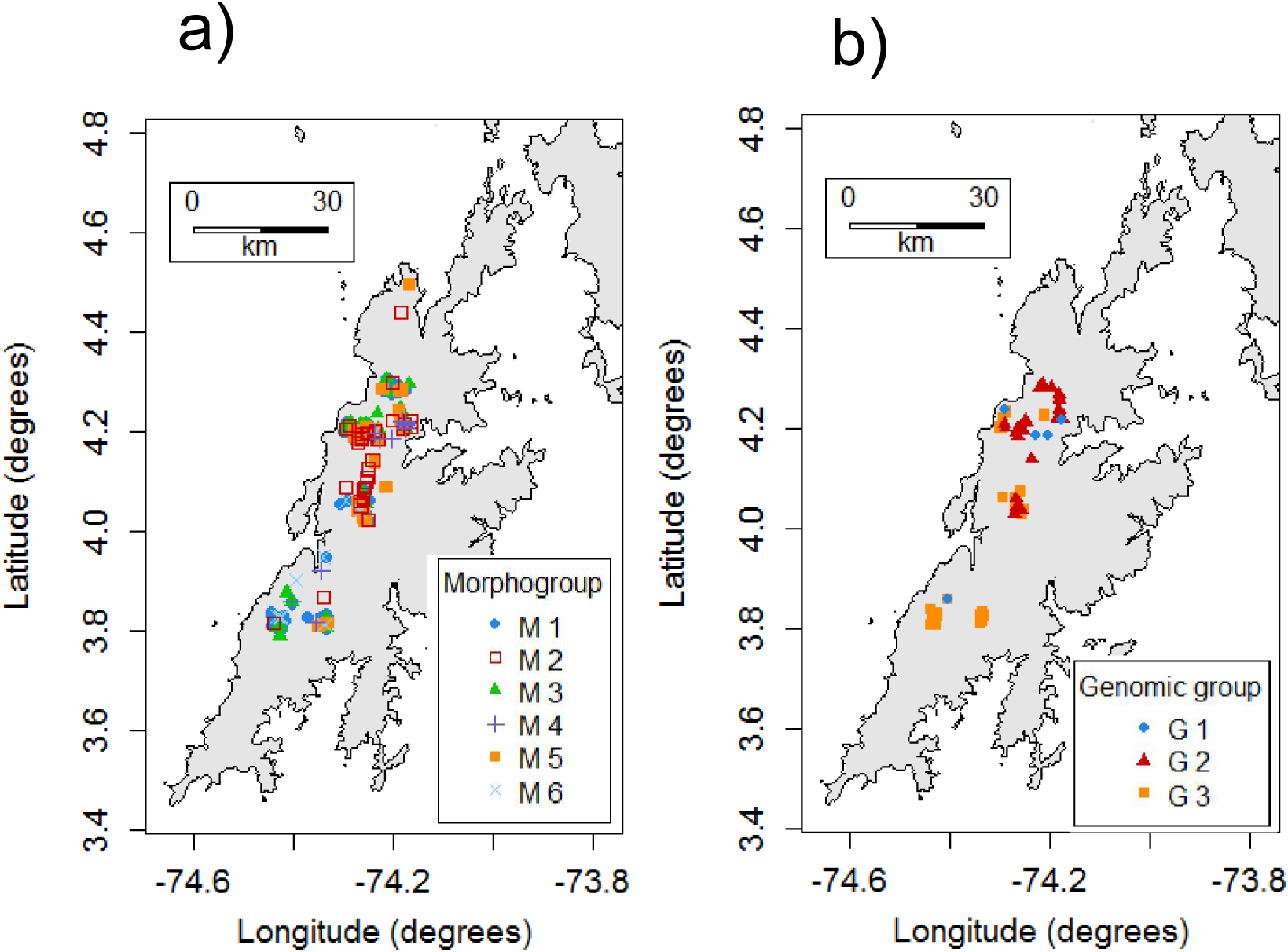
All pairs of **a)** phenotypic and **b)** genomic groups were sympatric, < 1 km apart. Contiguous areas above 3,000 m of elevation (gray, black contour) constitute the paramo de Sumapaz, separated from the paramo de Chingaza in the northeast. Specimen collection localities are shown as symbols according to phenotypic and genomic groups, as in Figures 5 and 6, respectively.

## DISCUSSION

Species are often regarded as basic units of biodiversity, but few empirical studies seem to directly address two key issues in a long-standing debate about the nature of species (Coyne and Orr 2004; Barraclough and Humphreys 2015; Barraclough 2019). One of these issues is whether organisms form sympatric groups, distinct in terms concordant phenotypic properties and genome-wide allele frequencies (Fig. 1). The other issue is the degree to which taxonomic classification at the species level (i.e., species taxa) accurately reflects distinct groups of individual organisms or constitute arbitrary divisions of biological diversity. Here we addressed both issues by studying plants of the Andean genus *Espeletia* (Asteraceae), based on geographically dense sampling of organisms in the páramo de Sumapaz, and the use of formal approaches to *de novo* inference of groups of organisms in terms of phenotype and genome-wide allele frequencies, as well as to measuring concordance between groups defined by phenotypes, genome-wide allele frequencies and species taxa.

To our surprise, we found that species taxa delimited in the most recent monograph of the group (Cuatrecasas 2013) were mostly empty in the sense that they contained only one of the 307 *Espeletia* specimens from Sumapaz in our analyses. In other words, we did not find the species taxa of Cuatrecasas (2013) in nature. However, analysis of data on 13 morphological characters did reveal fairly distinct, sympatric phenotypic groups among the *Espeletia* from Sumapaz. On the other hand, analysis of genome-wide allele frequencies indicated somewhat indistinct, sympatric groups with high levels of admixture among the *Espeletia* from Sumapaz. These phenotypic and genomic groups were only weakly concordant. Therefore, among the various hypotheses regarding the nature of species, our results seem most consistent with that in Figure 1c and d. The *Espeletia* from Sumapaz seem to form discordant sets of distinct phenotypes and less distinct groups of genome-wide allele frequencies, suggesting that phenotypes and genome-wide allele frequencies reflect contrasting evolutionary histories incorporated into a single species. Next, we explore implications of these results and potential caveats in our analyses.

Normal mixture models (NMMs) revealed distinct, sympatric morphological groups, which suggests the existence of six, non-cryptic *Espeletia* species in Sumapaz, thus ruling out scenarios depicted in Figure 1a and e. This conclusion is based on the assumptions needed to apply NMMs to species delimitation based on phenotypic characters (Cadena et al. 2018), including polygenic inheritance. We are not aware of studies of the genetic architecture of morphological characters in *Espeletia*. However, studies of quantitative trait loci (QTL) in confamilial species suggests that morphological characters (including some in Table 1) are determined by QTL of small effects (relative to the variance of inter-specific backcross populations, Lexer et al. 2005), as expected for polygenic inheritance. Two additional assumptions implicit in the application of NMMs to our context are that morphological groups do not reflect ontogenetic variation or phenotypic plasticity (Cadena et al. 2018). Given that all specimens included in the analysis were adult reproductive plants, we expect only negligible ontogenetic effects. Also, morphological groups seemed to share environments, at least broadly defined, as suggested by extensive sympatry (Fig. 6). In sum, the assumptions required by the application of NMMs seem to be reasonable for our study system.

We found evidence of three sympatric groups of *Espeletia* specimens from Sumapaz, characterized by different allelic frequencies across the genome. This finding is based on ancestry models that assume Hardy-Weinberg equilibrium and linkage equilibrium within groups (Pritchard et al. 2000). These models are often taken as a useful but rough approximation to reality, and it may be unwise to place much emphasis on any single value for the number of genomic groups (in our case 3). It may be more useful to consider the entire distribution of posterior probabilities for the number of genomic groups (Verity and Nichols 2016). Nonetheless, in our case the posterior distribution for the number of groups takes a value close to one for 3 groups and close to zero for all other values (Fig. 4b). Thus, it seems reasonable to conclude that there is strong support for *the* existence of three genomic groups. However, in contrast to the morphological groups, these genomic groups seemed indistinct and characterized by high levels of admixture (Fig. 6a). In fact, admixture values near 50% for several specimens in our sample suggest that F1 hybrids among the three genomic groups may be common.

The weak concordance between phenotypic and genomic groups suggests that the nature of *Espeletia* species in Sumapaz resembles that depicted in Figure 1d. However, the genomic groups are poorly differentiated (Fig. 6a). Therefore, the nature of *Espeletia* species in Sumapaz may be more similar to the hypothesis depicted in Figure 1c, a syngameon. In both cases (Fig. 1 c and d) phenotypic and genomic distinctiveness reflect contrasting evolutionary histories (Doyle 1997; Maddison 1997; Szllosi et al. 2015). This lack of agreement may be due to the fact that a general scan of the allele frequencies across the genome may not represent small and atypical parts of the genome that determine phenotypic distinction. For instance, in a syngameon frequent hybridization and introgression may homogenize large sections of the genome of different species that, nonetheless, remain phenotypically distinct due to selection on relatively few loci, as seems to be the case in some oaks (Van Valen 1976; Templeton 1989, Hipp et al. 2019). This idea is consistent with high admixture levels in our sample of *Espeletia* from Sumapaz, suggesting that F1 hybrids among genomic groups are common (Fig. 6a), and with previous studies of *Espeletia* that inferred hybridization between distantly related taxa (Pouchon et al. 2018), measured high crossability between species taxa (Berry et al. 1988) and observed putative hybrids between several pairs of species taxa (Cuatrecasas 2013; Diazgranados and Barber 2017).

We did not find the species taxa described in the most recent monograph of *Espeletia* (Cuatrecasas 2013) for the páramo de Sumapaz. This result suggests that species taxa are not even arbitrary divisions of biological diversity, as proposed by botanists (Levin 1979; Raven 1986; Bachmann 1998) and suggested by Darwin (1859). Far more disconcerting, species taxa in *Espeletia* appear to largely miss biological diversity, delineating mostly empty phenotypic space. We wonder if this result is unique to our study taxa and region. We note, however, that formal, specimen-based exploration of descriptions of species taxa in multidimensional phenotypic space is rarely carried out. This is no small matter, given that species are considered basic units of biodiversity in ecology, evolution, biogeography and conservation biology (Coyne and Orr 2004; Richards 2010; Barraclough and Humphreys 2015; Sigwart 2018). In this study we provided an example of how such exploration may be accomplished. In any case, our findings regarding species taxa strongly suggest the need to revise species limits in *Espeletia*, in addition to the taxonomic changes above the species level suggested by Pouchon et al. (2018).

## ACKNOWLEDGMENTS

This study is part of a long-term project on the effect of global change on *Espeletia*, designed and coordinated by Parques Nacionales Naturales de Colombia (PNN), the Jardín Botánico Bogotá José Celestino Mutis (JBB) and the Missouri Botanical Garden (MBG). The herbarium JBB provided key support and infrastructure to process specimens and, together with PNN, to conduct field work in the páramo de Sumapaz. The herbarium of the Universidad Nacional (COL) welcomed us into their collection. We thank the campesino communities of Sumapaz and the Ballón de Alta Montaña No. 1 for hosting and supporting us during field work. The data in this study was gathered with significant field and herbarium assistance from Erika Daniela Camacho, Alvaro Andrés Mariño, David Andrés Quinche, David Julián Forero and Juan Diego Martínez. Betsy Viviana Rodríguez provided important insight during herbarium work. Special thanks to Cristian Tibabisco, Erika Benavides, Mailo Benavides, Moisés Penagos, Yadira Baquero and Paola Medina for support during field work. This study was funded by a National Geographic grant (number WW-147R-17) to Iván Jiménez (MBG), Carlos A. Lora (PNN) and César Marín (JBB), and by a grant from Fondo de Investigaciones de la Facultad de Ciencias of the Universidad de los Andes to Yam M. Pineda. SNP genotyping was funded by the Vetenskapsrådet (VR) and Kungliga Vetenskapsakademien (KVA) grants to Andrés J. Cortés, with grant numbers 4.J1-2016-00418 and BS2017-0036, respectively.

## LITERATURE CITED

Adler P.B., HilleRislambers J., Levine J.M. 2007. A niche for neutrality. Ecol. Lett. 10:95–104.

Agresti A. 2002. Categorical data analysis. Hoboken (NJ): John Wiley & Sons.

Allmon W.D. 2016. Studying species in the fossil record?: a review and recommendations for a more unified approach. In: Allmon W.D., Yacobucci M.M., editors. Species and speciation in the fossil record. Chicago (IL): University of Chicago Press. p. 59–120.

Arnold M.L. 2015. Divergence with genetic exchange. OUP Oxford. 272 p.

Bachmann K. 1998. Species as units of diversity: an outdated concept. Theory Biosci. 117:213–230.

Barraclough T.G. 2019. The evolutionary biology of species. Imperial Collage London, UK. Oxford Series in Ecology and Evolution. Oxford University Press. 284 p.

Barraclough T.G., Humphreys A.M. 2015. The evolutionary reality of species and higher taxa in plants: A survey of post-modern opinion and evidence. New Phytol. 207:291–296.

Berry P., Beaujon S., Calvo R. 1988. La hibridización en la evolución de los frailejones (Espeletia, Asteraceae). Ecotropicos. 1:11–24.

Berry P.E., Calvo R.N. 1989. Wind Pollination, Self-Incompatibility, and Altitudinal Shifts in Pollination Systems in the High Andean genus Espeletia (Asteraceae). Am. J. Bot. 76:1602–1614.

Bickford D., Lohman D.J., Sodhi N.S., Ng P.K.L., Meier R., Winker K., Ingram K.K., Das I. 2007. Cryptic species as a window on diversity and conservation. Trends Ecol. Evol.: 22, 148–155.

Cadena C.D., Zapata F., Jiménez I. 2018. Issues and perspectives in species delimitation using phenotypic data: Atlantean evolution in Darwin’s Finches. Syst. Biol. 67:181–194.

Cavender-Bares J., Pahlich A. 2009. Molecular, morphological, and ecological niche differentiation of sympatric sister oak species, Quercus virginiana and Q. geminata (Fagaceae). Am. J. Bot. 96:1690–1702.

Chesson P. 2000. of S Pecies D Iversity. Annu. Rev. Ecol. Syst. 31:343–66.

Cortés A.J., Garzón L.N., Valencia J.B., Madriñán S. 2018. On the Causes of Rapid Diversification in the Páramos: Isolation by Ecology and Genomic Divergence in Espeletia. Front. Plant Sci. 9:1–17.

Coyne J.A., Orr H.A. 2004. Speciation. Sunderland (MA): Sinauer Associates. Cracraft. 545p.

Crisp M.D., Weston P.H. 1993. Geographic and Ontogenetic Variation in Morphology of Australian Waratahs (Telopea: Proteaceae). Syst. Biol. 42:49–76.

Cuatrecasas J. 2013. A Systematic Study of the Subtribe Espeletiinae (Heliantheae, Asteraceae). Volume 107; The New York Botanical Garden Press. 689 p.

Darwin C. 1859. The origin of species. Routledge. 502 p.

Diamond J.M. 1992. 6Horrible plant species. Nature. 360:627–628.

Diazgranados M. 2012. A nomenclator for the frailejones (Espeletiinae Cuatrec., Asteraceae). PhytoKeys. 16:1–52.

Diazgranados M., Barber J.C. 2017. Geography shapes the phylogeny of frailejones (Espeletiinae Cuatrec., Asteraceae): a remarkable example of recent rapid radiation in sky islands. PeerJ. 5:e2968.

Dobzhansky T. 1951. Genetics and the Origin of Species. 3rd edition, New York: Columbia Univ. Press.

Doyle J. 1997. Trees Within Trees?: Genes and Species, Molecules and Morphology. Syst. Biol. 46:537–553.

Ehrlich P.R., Raven P.H. 1969. Differentiation of populations published. Science (80-.). 165:1228–1232.

Elshire R.J., Glaubitz J.C., Sun Q., Poland J.A., Kawamoto K., Buckler E.S., Mitchell S.E. 2011. A robust, simple genotyping-by-sequencing (GBS) approach for high diversity species. PLoS One. 6:1–10.

Endersby J. 2009. Lumpers and splitters: Darwin, Hooker, and the search for order. Science (80-.). 326:1496–1499.

Ezard T.H., Pearson P.N., Purvis A. 2010. Algorithmic approaches to aid species’ delimitation in multidimensional morphospace. BMC Evol. Biol. 10:1–11.

Fagua J.C., Gonzalez V.H. 2007. Growth rates, reproductive phenology, and pollination ecology of Espeletia grandiflora (Asteraceae), a giant Andean caulescent rosette. Plant Biol. 9:127–135.

Fišer C., Robinson C.T., Malard F. 2018. Cryptic species as a window into the paradigm shift of the species concept. Mol. Ecol. 27:613–635.

Fraley C., Raftery A.E. 2002. Model-Based Clustering, Discriminant Analysis, and Density Estimation. J. Am. Stat. Assoc. 97:611–631.

Futuyma D.J., Mayer G.C. 1980. Non-Allopatric Speciation in Animals. Syst. Zool. 29:254–271.

Gallego Maya A., Bonilla Gómez M. 2016. Caracterización de micrositios para el establecimiento de plántulas de Espeletia uribei (Asteraceae). Acta Biológica Colomb. 21:387–398.

Glaubitz J.C., Casstevens T.M., Lu F., Harriman J., Elshire R.J., Sun Q., Buckler E.S. 2014. TASSEL-GBS: A high capacity genotyping by sequencing analysis pipeline. PLoS One. 9:e90346. doi:10.1371/journal.pone.0090346.

Gould S. 2002. The Structure of Evolutionary Theory. Harvard University Press.

Goulson D., Stout J. 2001. Homing ability of the bumblebee Bombus terrestris (Hymenoptera: Apidae). Sciences (New. York). 15:199–207.

Grant V. 1957. The plant species in theory and practice. Species Probl. (ed. Mayr, E.). Am. Assoc. Adv. Sci. Washint.38–90.

Grant V. 1971. Plant Speciation. Verne Grant. Columbia University Press, New York, 1971. xii, 436 pp., illus. $15. Columbia Univ. Press. New York. Xii:436.

Greenleaf S.S., Williams N.M., Winfree R., Kremen C. 2007. Bee foraging ranges and their relationship to body size. Oecologia. 153:589–596.

Hey J. 2001. Genes, Categories and Species: The evolutionary and cognitive causes of the species problem. Oxford University Press.

Hey J., Waples R.S., Arnold M.L., Butlin R.K., Harrison R.G. 2003. Understanding and confronting species uncertainty in biology and conservation. Trends Ecol. Evol. 18:597–603.

Hipp A., Whittemore A., Garner M., Guichoux E., Hahn M., Fitzek E., Cavender-bares J., Gugger P.F., Manos P.S., Pearse I.S., Cannon C.H. 2019. Conserved DNA polymorphisms distinguish species in the eastern North American white oak syngameon: Insights from an 80-SNP oak DNA genotyping toolkit. bioRxiv. 602573.

Korshunova T., Picton B., Furfaro G., Mariottini P., Pontes M., Prkić J., Fletcher K., Malmberg K., Lundin K., Martynov A. 2019. Multilevel fine-scale diversity challenges the ‘cryptic species’ concept. Sci. Rep. 9:1–23.

Lawson D.J., van Dorp L., Falush D. 2018. A tutorial on how not to over-interpret structure and admixture bar plots. Nat. Commun. 9:1–11.

Levin D.A. 1979. The nature of plant species: Science. 204:381–384.

Lexer C., Rosenthal D.M., Raymond O., Donovan L.A., Rieseberg L.H. 2005. Genetics of species differences in the wild annual sunflowers, Helianthus annuus and H. petiolaris. Genetics. 169:2225–2239.

Lotsy J.P. 1931. On the species of the taxonomist in its relation to evolution. Genetica. 13:1–16.

Maddison W.P. 1997. Gene trees in species trees. Syst. Biol. 46:523–536.

Madriñán S., Cortés A.J., Richardson J.E. 2013. Páramo is the world’s fastest evolving and coolest biodiversity hotspot. Front. Genet. 4:1–7.

Mallet J. 2007. Concepts of Species. Encycl. Biodivers.1–15.

Mallet J. 2008. Hybridization, ecological races and the nature of species: empirical evidence for the ease of speciation. Proc. R. Soc. 363:2971–2986.

Mallet J. 2013. Species, Concept of. Encycl. Biodivers. 679:9–15.

Mallet J., Meyer A., Nosil P., Feder J.L. 2009. Space, sympatry and speciation. J. Evol. Biol. 22:2332–2341.

Maugis C., Celeux G., Martin-Magniette M.L. 2009. Variable selection for clustering with gaussian mixture models. Biometrics. 65:701–709.

Mayr E. 1947. Ecological Factors in Speciation. Evolution (N. Y). 1:263–288.

Mayr E. 1963. Animal Species and Evolution. Cambridge. The Belknap Press of Harvard University Press. 797 p.

Mayr E. 1982. The Growth of Biological Thought Diversity, Evolution, and Inheritance. Harvard University Press. 974 p.

Mayr E. 1992. A local flora and the biological species concept. Am. J. Bot. 79:222–238.

McLachlan G., Peel D. 2000. Finite Mixture Models. A wile-Interscience Publication, John wiley & Sons, inc. New York: Hoboken (NJ): John Wiley and Sons (Series in probability and statistics). 438 p.

McLachlan G.J. 2004. Discriminant analysis and statistical pattern recognition. Hoboken (NJ): John Wiley and Sons (Series in probability and statistics). 526 p.

Monasterio M., Sarmiento L. 1991. Adaptive Radiation of Espeletia in Cold Andean Tropics. Tree Trends Ecol. Evol. 6:387–391.

Pearson R. 2016. Association Analysis for Categorical Variables. Package ‘GoodmanKruskal.’ CRAN.:https://cran.r-project.org/web/packages/GoodmanKru.

Pope N.S., Jha S. 2018. Seasonal Food Scarcity Prompts Long-Distance Foraging by a Wild Social Bee. Am. Nat. 191:45–57.

Pouchon C., Fernández A., Nassar J.M., Boyer F., Aubert S., Lavergne S., Mavárez J. 2018. Phylogenomic analysis of the explosive adaptive radiation of the Espeletia complex (Asteraceae) in the tropical Andes. Syst. Biol. 67:1041–1060.

Pritchard J.K., Stephens M., Donnelly P. 2000. Inference of Population Structure Using Multilocus Genotype Data. Genet. Soc. Am. 155:945–959.

De Queiroz K. 1999. The general lineage concept of species and the defining properties of the species category. Species, New interdisciplinary essays. p. 49–89.

De Queiroz K. 2007. Species concepts and species delimitation. Syst. Biol. 56:879–886.

De Queiroz K. 2011. Branches in the lines of descent: Charles Darwin and the evolution of the species concept. Biol. J. Linn. Soc. 103:19–35.

Queiroz K.D.E., Good A. 1997. Phenetic clustering in biology: a critique. Q. Rev. Biol. 72:3–30.

Raftery A.E., Dean N. 2006. Variable selection for model-based clustering. J. Am. Stat. Assoc. 101:168–178.

Ramsay P.M., Oxley E.R. 1997. The growth form composition of plant communities in the Ecuadorian páramos. Plant Ecol. 131:173–192.

Rauscher J.T. 2002. Molecular Phylogenetics of the Espeletia Complex (Asteraceae): Evidence from nrDNA ITS Sequences on the Closest Relatives of an Andean Adaptive Radiation. Am. J. Bot. 89:1074–1084.

Raven P.H. 1986. Modern aspects of the biological species in plants. eds Kunio Iwatsuki, Peter H. Raven, and Walter J. Bock. University of Tokyo Press.

Richards R.A. 2010. The Species Problem A philosophical Analysis. Published in the United States of America by Cambridge University Press, New York. 347 p.

Rieseberg L.H., Burke J.M. 2001. A genic view of species integration. 14:883–886.

Rieseberg L.H., Wood T.E., Baack E.J. 2006. The nature of plant species. Nature. 440:524–527.

Schindelin C.A., Rasband W.S., Eliceiri K.W. 2012. NIH Image to ImageJ: 25 years of image analysis. Nat. Methods. 9:671.

Schwartz G. 1978. Estimating the Dimension of a Model. Ann. Stat. 6:461–464.

Scrucca L., Fop M., Murphy T.B., Raftery A.E. 2016. mclust 5: Clustering, Classification and Density Estimation Using Gaussian Finite Mixture Models. R J. 8:289–317.

Scrucca L., Raftery A.E. 2015. Improved initialisation of model-based clustering using Gaussian hierarchical partitions. Adv. data Anal. Classif. 9:447–460.

Scrucca L., Raftery A.E. 2018. clustvarsel: A Package Implementing Variable Selection for Model-based Clustering in R. J. Stat. Software, Artic. 84.

Sheth S.N., Lohmann L.G., Distler T., Jiménez I. 2012. Understanding bias in geographic range size estimates. Glob. Ecol. Biogeogr. 21:732–742.

Sigwart J.D. 2018. What Species Mean A Users Guide to the Units of Biodiversity. Kipling Will (University of California, Berkeley), CRC Press Taylor & Francis Group: Charles R. Crumly, CRC Press/Taylor and Francis.

Stamos D.N. 2007. Darwin and the Nature of Species. State Univ. New York Press. Albany.:1–21.

ter Steege H., Haripersaud P.P., Bánki O.S., Schieving F. 2011. A model of botanical collectors’ behavior in the field: never the same species twice. Am. J. Bot. 98:31–7.

Stevens P.F. 1997. J.D. Hooker, George Bentham, Asa gray and ferdinand mueller on species limits in theory and practice: A mid-nineteenth-century debate and its repercussions. Hist. Rec. Aust. Sci. 11:345–370.

Struck T.H., Feder J.L., Bendiksby M., Birkeland S., Cerca J., Gusarov V.I., Kistenich S., Larsson K.H., Liow L.H., Nowak M.D., Stedje B., Bachmann L., Dimitrov D. 2018. Finding Evolutionary Processes Hidden in Cryptic Species. Trends Ecol. Evol. 33:153–163.

Szllosi G.J., Tannier E., Daubin V., Boussau B. 2015. The inference of gene trees with species trees. Syst. Biol. 64:e42–e62.

Templeton A.R. 1989. The meaning of species and speciation: a genetic perspective. Speciation and its consequences. D. Otte and J. A. Endler, eds. Sinauer Associates, Sunderland, Massachusetts.: p. 3–27.

Templeton A.R. 2006. Population Genetics and Microevolutionary Theory. A John Wiley & Sons., Inc., Publication.

Van Valen L. 1976. Ecological species, multispecies and oaks. Taxon. 25:233–239.

Verity R., Nichols R.A. 2016. Estimating the number of subpopulations (K) in structured populations. Genetics. 203:1827–1835.

Wilkins J.S. 2018. Species: The Evolution of the Idea. CRC Press.

Wu C.-I. 2001. The genic view of the process of speciation. Evol.Biol. 14:851–865.

