## Supplementary Material in Appendix 2 for "The Nature of *Espeletia* Species"

###################################################################################################################
###################################################################################################################
###################################################################################################################
#
# INTRODUCTION
# This script is part of the supplementary materials for Pineda et al. (The Nature of Espeletia
# Species). It performs species delimitation analyses based onmphenotypic data taken on specimens
# of the genus Espeletia (Asteraceae) form the p�ramo de Sumapaz in the Eastern Andes of Colombia. 
# 
# PURPOSE OF THIS SCRIPT
# The script creates a dataframe with mean values of 13 phenotypic characters for each Espeletia specimen
# measured, and then performs analyses to identify phenotypic groups of specimens based on normal mixture
# models.
#
# DATA FILES REQUIRED TO RUN THIS SCRIPT
# Data file "Pineda_etal_BioRxiv_Appendix1.csv", with measurements of pheotypic characters for each Espeletia
# specimen. This file is available as Supplementary Material in Appendix 1.
# 
# TABLE OF CONTENTS
# 1) Preliminaries: load packages, read data, calculate mean values per trait for each specimen.
# 2) Examine the distribution of phenotypic trait values, edit data, transform and rotate data with 
#    principal component analysis.
# 3) Variable selection for normal mixture models.
# 4) Fit normal mixture models.
# 5) Examine phenotypic groups in the best normal mixture model.
#
###################################################################################################################
###################################################################################################################
###################################################################################################################


###################################################################################################################
###################################################################################################################
# 1) Preliminaries: load packages, read data, calculate mean values per trait for each specimen.
###################################################################################################################
###################################################################################################################

###################################################################################################################
# 1.1) Load packages.

library(clustvarsel) #package to perform variable selection for normal mixture models
library(mclust) #package to fit normal mixture models

###################################################################################################################
# 1.2) Read data.

#set working directory
#setwd("C:/_transfer/Review/MelissaPineda/Data_Melissa") #Ivan's working directory Lehmann
#setwd("C:/_transfer/Proposals/Espeletia/TesisMelissa/Data") #Ivan's working directory Waterman
setwd("~/Desktop/Tesis_Frailejones/") #Melissa's laptop

#read phenotypic data
phenodata <- read.table("phenodata_2019Jul25.csv", header=T, sep=",")

#examine phenotypic data
summary(phenodata)
head(phenodata)
dim(phenodata)

###################################################################################################################
# 1.3) For each specimen and each phenotypic character, calculate mean values across multiple structures measured.

colnames(phenodata)[9:11]
Length.Lamina <- rowMeans(phenodata[,9:11], na.rm=T)
#summary(Length.Lamina)
#sum(is.nan(Length.Lamina))
Length.Lamina[is.nan(Length.Lamina)] <- NA

colnames(phenodata)[12:14]
Width.Lamina <- rowMeans(phenodata[,12:14], na.rm=T)
Width.Lamina[is.nan(Width.Lamina)] <- NA

colnames(phenodata)[15:17]
Length.Synflorescence.Axis <- rowMeans(phenodata[,15:17], na.rm=T)
Length.Synflorescence.Axis[is.nan(Length.Synflorescence.Axis)] <- NA

colnames(phenodata)[18:20]
Perimeter.Synflorescence.Axis <- rowMeans(phenodata[,18:20], na.rm=T)
Perimeter.Synflorescence.Axis[is.nan(Perimeter.Synflorescence.Axis)] <- NA

colnames(phenodata)[21:23]
Number.Pairs.Sterile.Leaves.Per.Synflorescence <- rowMeans(phenodata[,21:23], na.rm=T)
Number.Pairs.Sterile.Leaves.Per.Synflorescence[is.nan(Number.Pairs.Sterile.Leaves.Per.Synflorescence)] <- NA

colnames(phenodata)[24:26]
Number.Capitula.Per.Synflorescence <- rowMeans(phenodata[,24:26], na.rm=T)
Number.Capitula.Per.Synflorescence[is.nan(Number.Capitula.Per.Synflorescence)] <- NA

colnames(phenodata)[27:32]
Length.Cyme.Peduncle <- rowMeans(phenodata[,27:32], na.rm=T)
Length.Cyme.Peduncle[is.nan(Length.Cyme.Peduncle)] <- NA

colnames(phenodata)[33:40]
Diameter.Capitulum <- rowMeans(phenodata[,33:40], na.rm=T)
Diameter.Capitulum[is.nan(Diameter.Capitulum)] <- NA

colnames(phenodata)[41:43]
Length.Sterile.Phyllary <- rowMeans(phenodata[,41:43], na.rm=T)
Length.Sterile.Phyllary[is.nan(Length.Sterile.Phyllary)] <- NA

colnames(phenodata)[44:46]
Width.Sterile.Phyllary <- rowMeans(phenodata[,44:46], na.rm=T)
Width.Sterile.Phyllary[is.nan(Width.Sterile.Phyllary)] <- NA

colnames(phenodata)[47:49]
Length.Ray.Flower.Corolla <- rowMeans(phenodata[,47:49], na.rm=T)
Length.Ray.Flower.Corolla[is.nan(Length.Ray.Flower.Corolla)] <- NA

colnames(phenodata)[50:52]
Length.Disc.Flower.Corolla <- rowMeans(phenodata[,50:52], na.rm=T)
Length.Disc.Flower.Corolla[is.nan(Length.Disc.Flower.Corolla)] <- NA

colnames(phenodata)[53:55]
Length.Disc.Flower.Tube <- rowMeans(phenodata[,53:55], na.rm=T)
Length.Disc.Flower.Tube[is.nan(Length.Disc.Flower.Tube)] <- NA

###################################################################################################################
# 1.4) Create a dataframe with mean values for each specimen and each phenotypic character.

mean.phenodata <- data.frame(phenodata[,1:8], Length.Lamina, Width.Lamina, Length.Synflorescence.Axis,
                             Perimeter.Synflorescence.Axis, Number.Pairs.Sterile.Leaves.Per.Synflorescence, Number.Capitula.Per.Synflorescence,
                             Length.Cyme.Peduncle, Diameter.Capitulum, Length.Sterile.Phyllary, Width.Sterile.Phyllary,
                             Length.Ray.Flower.Corolla, Length.Disc.Flower.Corolla, Length.Disc.Flower.Tube, phenodata[,56:59]) 
head(mean.phenodata)
summary(mean.phenodata)
dim(mean.phenodata)


###################################################################################################################
###################################################################################################################
# 2) Examine the distribution of each phenotypic trait, edit data, transform and rotate data with 
#    principal component analysis.
###################################################################################################################
###################################################################################################################

###################################################################################################################
# 2.1) Graphically examine the distribution of each phenotypic trait in linear and logarithmic scales.

colnames(mean.phenodata) #these are column names, showing the names of phenotypic traits 
trait.x <-21 #select a phenotypic trait by selecting a column
colnames(mean.phenodata)[trait.x] #check the name
#trait distribution in linear scale
hist(mean.phenodata[,trait.x], breaks=100, xlab=colnames(mean.phenodata)[trait.x], main="", col="gray80")
summary(mean.phenodata[,trait.x])
#trait distribution in lograrithmic scale
hist(log(mean.phenodata[,trait.x]), breaks=100, xlab=colnames(mean.phenodata)[trait.x], main="", col="gray80")
summary(log(mean.phenodata[,trait.x]))
#trait distribution in lograrithmic scale after adding one, this may be useful when there are zeros in the raw data 
hist(log(mean.phenodata[,trait.x]+1), breaks=100, xlab=colnames(mean.phenodata)[trait.x], main="", col="gray80")
summary(log(mean.phenodata[,trait.x]+1))

#graphically examine bivariate trait relationships,
#define the two traits to examine (select two columns of data frame mean.phenodata)
colnames(mean.phenodata) #these are column names, showing the names of phenotypic traits 
trait.x <- 9
trait.y <- 10
#check the names of selected traits
colnames(mean.phenodata)[trait.x]
summary(mean.phenodata[,trait.x])
colnames(mean.phenodata)[trait.y]
summary(mean.phenodata[,trait.y])
#plot the bivariate relationship
plot(mean.phenodata[,trait.x], mean.phenodata[,trait.y], xlab=names(mean.phenodata)[trait.x], 
	ylab=names(mean.phenodata)[trait.y], cex.lab=1.5, cex.axis=1.5)
abline(0,1)

###################################################################################################################
# 2.2) Select a subset of phenotypic traits to include in the analysis. This selection of traits is ideally
# done according to a priori ideas about the importance of characters to distinguish species, based on the
# Espeletiinae monograph (Cuatrecasas, 2013). 

#select a subset of phenotypic traits
colnames(mean.phenodata) #these are column names, showing the names of phenotypic traits
mean.phenodata.selected <- mean.phenodata[,c(9,10,11,12,13,14,15,16,17,18,19,20,21)]

#examine the results
dim(mean.phenodata.selected)
summary(mean.phenodata.selected)
head(mean.phenodata.selected)

###################################################################################################################
# 2.3) Remove specimens that have NA values for any of the phenotypic traits selected in the previous section.

#determine which specimens (i.e., rows in the data frame) have
#NA values; those specimens will have to be excluded from the analysis.
rows.with.na <- unique(which(is.na(mean.phenodata.selected), arr.ind = T)[,1])
rows.with.na #these are the specimens (i.e., rows) with NA values
length(rows.with.na) #this is the number  specimens with NA values
#run the following lines only if there are specimens with NA values
mean.phenodata.selected <- mean.phenodata.selected[-rows.with.na,]
dim(mean.phenodata.selected) #after removing NA values, 307 specimens remain in the data frame
class(mean.phenodata.selected)
summary(mean.phenodata.selected)
head(mean.phenodata.selected, 5)
rownames(mean.phenodata.selected)

###################################################################################################################
# 2.4) Transform data.

#log-transform continuous traits, given continuous phenotypic traits often approximately follow a log-normal
#distribution, be sure to add 1 to traits whose values are left-bounded by zero.

mean.phenodata.selected.log <- log(data.frame(mean.phenodata.selected[,1:4], mean.phenodata.selected[,5]+1, 
	mean.phenodata.selected[,6:13]))
head(mean.phenodata.selected.log)

#edit column names
colnames(mean.phenodata.selected.log) <- paste("log", colnames(mean.phenodata.selected))
colnames(mean.phenodata.selected.log)[5] <- paste("log", paste(colnames(mean.phenodata.selected)[5], "+1", sep=""))
class(mean.phenodata.selected.log)
summary(mean.phenodata.selected.log)

###################################################################################################################
# 2.5) Perform principal component analysis (PCA) on the covariance matrix of the log-transformed continuous
# traits.

#perform PCA using the covariance matrix
mean.phenodata.selected.log.pca <- prcomp(mean.phenodata.selected.log, center = T, scale. = F) #PCA using the covariance matrix
mean.phenodata.selected.log.pca

#examine PCA results
attributes(mean.phenodata.selected.log.pca) #attributes of the R object (class "prcomp") containing the PCA results 
mean.phenodata.selected.log.pca$scale
mean.phenodata.selected.log.pca$center
summary(mean.phenodata.selected.log.pca) #variance explained by each principal component
summary(mean.phenodata.selected.log.pca$x) #summary of the principal components 
mean.phenodata.selected.log.pca$rotation #coefficients (or "loadings") of each trait on each principal component

#examine univariate distributions of principal components:
#select a principal component (select a column of data frame mean.phenodata.con.log.pca$x)
colnames(mean.phenodata.selected.log.pca$x) #column names corresponding to principal components
PCA.x <- 1
#check the name
colnames(mean.phenodata.selected.log.pca$x)[PCA.x]
#examine histogram
hist(mean.phenodata.selected.log.pca$x[,PCA.x], breaks=100, xlab=colnames(mean.phenodata.selected.log.pca$x)[PCA.x],
	ylab="Specimens", main="", cex.lab=1.5, cex.axis=1.5)

##examine bivariate relationships between principal components 
#select two principal components to examine (select two columns from mean.phenodata.selected.log.pca$x)
colnames(mean.phenodata.selected.log.pca$x) #column names corresponding to principal components
PCA.x <- 1
PCA.y <- 2
#check their names
dimnames(mean.phenodata.selected.log.pca$x)[[2]][PCA.x]
dimnames(mean.phenodata.selected.log.pca$x)[[2]][PCA.y]
#plot the bivariate relationship
plot(mean.phenodata.selected.log.pca$x[,PCA.x], mean.phenodata.selected.log.pca$x[,PCA.y],
	xlab=dimnames(mean.phenodata.selected.log.pca$x)[[2]][PCA.x], ylab=dimnames(mean.phenodata.selected.log.pca$x)[[2]][PCA.y],
	cex.lab=1.5, cex.axis=1.5)


###################################################################################################################
###################################################################################################################
# 3) Variable selection for normal mixture models.
###################################################################################################################
###################################################################################################################

###################################################################################################################
# 3.1) Backward variable selection using PCA on the covariace matrix of the log-transformed data.

#perform backward variable selection for different initialization approaches, as defined with argument "hcUse":
#check and set Mclust options as needed (see help for function "Mclust")
mclust.options()

mclust.options(hcUse="PCS") 
mean.phenodata.selected.log.pca.varsel.back <- clustvarsel(mean.phenodata.selected.log.pca$x, G=1:10, search=c("greedy"), direction = c("backward"))
#examine results
attributes(mean.phenodata.selected.log.pca.varsel.back)
summary(mean.phenodata.selected.log.pca.varsel.back)
names(mean.phenodata.selected.log.pca.varsel.back$subset) 
mean.phenodata.selected.log.pca.varsel.back$steps.info
mean.phenodata.selected.log.pca.varsel.back$search
mean.phenodata.selected.log.pca.varsel.back$direction
#these are the traits selected, in the order they were selected: 1,2,3,4,5,6,7,8,9,10,11,12

mclust.options(hcUse="VARS")
mean.phenodata.selected.log.pca.varsel.back <- clustvarsel(mean.phenodata.selected.log.pca$x, G=1:10, search=c("greedy"), direction = c("backward"))
#examine results
attributes(mean.phenodata.selected.log.pca.varsel.back)
summary(mean.phenodata.selected.log.pca.varsel.back)
names(mean.phenodata.selected.log.pca.varsel.back$subset) 
mean.phenodata.selected.log.pca.varsel.back$steps.info
mean.phenodata.selected.log.pca.varsel.back$search
mean.phenodata.selected.log.pca.varsel.back$direction
#these are the traits selected, in the order they were selected: 1,2,3,4,5,6,7,8,9,10,11,12

mclust.options(hcUse="STD")
mean.phenodata.selected.log.pca.varsel.back <- clustvarsel(mean.phenodata.selected.log.pca$x, G=1:10, search=c("greedy"), direction = c("backward"))
#examine results
attributes(mean.phenodata.selected.log.pca.varsel.back)
summary(mean.phenodata.selected.log.pca.varsel.back)
names(mean.phenodata.selected.log.pca.varsel.back$subset) 
mean.phenodata.selected.log.pca.varsel.back$steps.info
mean.phenodata.selected.log.pca.varsel.back$search
mean.phenodata.selected.log.pca.varsel.back$direction
#these are the traits selected, in the order they were selected: 1,2,3,4,5,6,7,8,9,10,11,12

mclust.options(hcUse="SPH")
mean.phenodata.selected.log.pca.varsel.back <- clustvarsel(mean.phenodata.selected.log.pca$x, G=1:10, search=c("greedy"), direction = c("backward"))
#examine results
attributes(mean.phenodata.selected.log.pca.varsel.back)
summary(mean.phenodata.selected.log.pca.varsel.back)
names(mean.phenodata.selected.log.pca.varsel.back$subset) 
mean.phenodata.selected.log.pca.varsel.back$steps.info
mean.phenodata.selected.log.pca.varsel.back$search
mean.phenodata.selected.log.pca.varsel.back$direction
#these are the traits selected, in the order they were selected: 1,2,3,4,5,6,7,8,9,10,11,12

mclust.options(hcUse="PCR")
mean.phenodata.selected.log.pca.varsel.back <- clustvarsel(mean.phenodata.selected.log.pca$x, G=1:10, search=c("greedy"), direction = c("backward"))
#examine results
attributes(mean.phenodata.selected.log.pca.varsel.back)
summary(mean.phenodata.selected.log.pca.varsel.back)
names(mean.phenodata.selected.log.pca.varsel.back$subset) 
mean.phenodata.selected.log.pca.varsel.back$steps.info
mean.phenodata.selected.log.pca.varsel.back$search
mean.phenodata.selected.log.pca.varsel.back$direction
#these are the traits selected, in the order they were selected: 1,2,3,4,5,6,7,8,9,10,11,12

mclust.options(hcUse="SVD")
mean.phenodata.selected.log.pca.varsel.back <- clustvarsel(mean.phenodata.selected.log.pca$x, G=1:10, search=c("greedy"), direction = c("backward"))
#examine results
attributes(mean.phenodata.selected.log.pca.varsel.back)
summary(mean.phenodata.selected.log.pca.varsel.back)
names(mean.phenodata.selected.log.pca.varsel.back$subset) 
mean.phenodata.selected.log.pca.varsel.back$steps.info
mean.phenodata.selected.log.pca.varsel.back$search
mean.phenodata.selected.log.pca.varsel.back$direction
#these are the traits selected, in the order they were selected: 1,2,3,4,5,6,7,8,9,10,11,12

###################################################################################################################
# 3.2) Forward variable selection using PCA on the covariace matrix of the log-transformed data.

#perform forward variable selection for different initialization approaches, as defined using argument "hcUse":
#check and set Mclust options as needed (see help for function "Mclust")
mclust.options()

mclust.options(hcUse="PCS")
mean.phenodata.selected.log.pca.varsel.for <- clustvarsel(mean.phenodata.selected.log.pca$x, G=1:10, search=c("greedy"), direction = c("forward"))
#examine results
attributes(mean.phenodata.selected.log.pca.varsel.for)
summary(mean.phenodata.selected.log.pca.varsel.for)
names(mean.phenodata.selected.log.pca.varsel.for$subset) 
mean.phenodata.selected.log.pca.varsel.for$steps.info
mean.phenodata.selected.log.pca.varsel.for$search
mean.phenodata.selected.log.pca.varsel.for$direction
#these are the traits selected, in the order they were selected: 1,3,2,6,5,4,7,8,9,10,12,11

mclust.options(hcUse="VARS")
mean.phenodata.selected.log.pca.varsel.for <- clustvarsel(mean.phenodata.selected.log.pca$x, G=1:10, search=c("greedy"), direction = c("forward"))
#examine results
attributes(mean.phenodata.selected.log.pca.varsel.for)
summary(mean.phenodata.selected.log.pca.varsel.for)
names(mean.phenodata.selected.log.pca.varsel.for$subset) 
mean.phenodata.selected.log.pca.varsel.for$steps.info
mean.phenodata.selected.log.pca.varsel.for$search
mean.phenodata.selected.log.pca.varsel.for$direction
#these are the traits selected, in the order they were selected: 1,3,2,6,5,4,7,8,9,10,12,11

mclust.options(hcUse="STD")
mean.phenodata.selected.log.pca.varsel.for <- clustvarsel(mean.phenodata.selected.log.pca$x, G=1:10, search=c("greedy"), direction = c("forward"))
#examine results
attributes(mean.phenodata.selected.log.pca.varsel.for)
summary(mean.phenodata.selected.log.pca.varsel.for)
names(mean.phenodata.selected.log.pca.varsel.for$subset) 
mean.phenodata.selected.log.pca.varsel.for$steps.info
mean.phenodata.selected.log.pca.varsel.for$search
mean.phenodata.selected.log.pca.varsel.for$direction
#these are the traits selected, in the order they were selected: 1,3,2,6,5,4,7,8,9,10,12,11

mclust.options(hcUse="SPH")
mean.phenodata.selected.log.pca.varsel.for <- clustvarsel(mean.phenodata.selected.log.pca$x, G=1:10, search=c("greedy"), direction = c("forward"))
#examine results
attributes(mean.phenodata.selected.log.pca.varsel.for)
summary(mean.phenodata.selected.log.pca.varsel.for)
names(mean.phenodata.selected.log.pca.varsel.for$subset) 
mean.phenodata.selected.log.pca.varsel.for$steps.info
mean.phenodata.selected.log.pca.varsel.for$search
mean.phenodata.selected.log.pca.varsel.for$direction
#these are the traits selected, in the order they were selected: 1,3,2,6,5,4,7,8,9,10,12,11

mclust.options(hcUse="PCR")
mean.phenodata.selected.log.pca.varsel.for <- clustvarsel(mean.phenodata.selected.log.pca$x, G=1:10, search=c("greedy"), direction = c("forward"))
#examine results
attributes(mean.phenodata.selected.log.pca.varsel.for)
summary(mean.phenodata.selected.log.pca.varsel.for)
names(mean.phenodata.selected.log.pca.varsel.for$subset) 
mean.phenodata.selected.log.pca.varsel.for$steps.info
mean.phenodata.selected.log.pca.varsel.for$search
mean.phenodata.selected.log.pca.varsel.for$direction
#these are the traits selected, in the order they were selected: 1,3,2,6,5,4,7,10,8,9,12,11

mclust.options(hcUse="SVD")
mean.phenodata.selected.log.pca.varsel.for <- clustvarsel(mean.phenodata.selected.log.pca$x, G=1:10, search=c("greedy"), direction = c("forward"))
#examine results
attributes(mean.phenodata.selected.log.pca.varsel.for)
summary(mean.phenodata.selected.log.pca.varsel.for)
names(mean.phenodata.selected.log.pca.varsel.for$subset) 
mean.phenodata.selected.log.pca.varsel.for$steps.info
mean.phenodata.selected.log.pca.varsel.for$search
mean.phenodata.selected.log.pca.varsel.for$direction
#these are the traits selected, in the order they were selected: 1,3,2,6,5,4,7,10,8,9,12,11


###################################################################################################################
###################################################################################################################
# 4) Fit normal mixture models.
###################################################################################################################
###################################################################################################################

###################################################################################################################
# 4.1) Select phenotypic variables (principal components) for inclusion in normal mixture models.

#based on results from backward and forward variable selection (sections 3.1 and 3.2)
data.for.GMM <- mean.phenodata.selected.log.pca$x[,c(1,2,3,4,5,6,7,8,9,10,11,12)]

###################################################################################################################
# 4.2) Fit normal mixture models using different initialization approaches, as defined using argument "hcUse"

#"PCS"
mclust.options(hcUse="PCS") 
Mcluster.phenodata <- Mclust(data.for.GMM)
#examine results
Mcluster.phenodata
summary(Mcluster.phenodata)
names(Mcluster.phenodata$classification)#the specimens included in the analysis
Mcluster.phenodata$classification #classification of specimens
Mcluster.phenodata$uncertainty #the uncertainty of the classification
attributes(Mcluster.phenodata)
#Mclust EVE (ellipsoidal, equal volume and orientation) model with 6 components:
# log-likelihood   n  df       BIC       ICL
# 306.4158       307  210 -589.8063 -603.7503
#Clustering table:
# 1   2  3  4  5  6 
#130  48 52 17 39 21
#graph morphogroups according to the best model
plot(Mcluster.phenodata, what="classification", dimens=c(1,2))
#graph empirical support for different models
plot(Mcluster.phenodata, what="BIC")
#save best Mclust model
#set working directory
#setwd("C:/_transfer/Review/MelissaPineda/Data_Melissa") #Ivan's working directory Lehmann
#setwd("C:/_transfer/Proposals/Espeletia/TesisMelissa/Data") #Ivan's working directory Waterman
#save(Mcluster.phenodata, file="Mcluster.phenodata_2020June09.RData")
#load("Mcluster.phenodata_2020June09.RData")

#"VARS"
mclust.options(hcUse="VARS")
Mcluster.phenodata <- Mclust(data.for.GMM)
#examine results
Mcluster.phenodata
summary(Mcluster.phenodata)
names(Mcluster.phenodata$classification)#the specimens included in the analysis
Mcluster.phenodata$classification #classification of specimens
Mcluster.phenodata$uncertainty #the uncertainty of the classification
attributes(Mcluster.phenodata)
#Mclust EVE (ellipsoidal, equal volume and orientation) model with 6 components: 
#log-likelihood   n  df       BIC       ICL
#306.4158         307 210 -589.8063 -603.7503
#Clustering table:
# 1   2   3   4   5   6 
#130  48  52  17  39  21 
#graph morphogroups according to the best model
plot(Mcluster.phenodata, what="classification", dimens=c(1,2))
#graph empirical support for different models
plot(Mcluster.phenodata, what="BIC")

#"STD"
mclust.options(hcUse="STD")
Mcluster.phenodata <- Mclust(data.for.GMM)
#examine results
Mcluster.phenodata
summary(Mcluster.phenodata)
names(Mcluster.phenodata$classification)#the specimens included in the analysis
Mcluster.phenodata$classification #classification of specimens
Mcluster.phenodata$uncertainty #the uncertainty of the classification
attributes(Mcluster.phenodata)
#Mclust VVE (ellipsoidal, equal orientation) model with 6 components: 
#log-likelihood   n  df       BIC       ICL
#    261.2505 	 307 215 -708.7712 -737.2176
#Clustering table:
# 1  2  3  4  5  6 
# 26 79 22 58 78 44 
#graph morphogroups according to the best model
plot(Mcluster.phenodata, what="classification", dimens=c(1,2))
#graph empirical support for different models
plot(Mcluster.phenodata, what="BIC")

#"SPH"
mclust.options(hcUse="SPH")
Mcluster.phenodata <- Mclust(data.for.GMM)
#examine results
Mcluster.phenodata
summary(Mcluster.phenodata)
names(Mcluster.phenodata$classification)#the specimens included in the analysis
Mcluster.phenodata$classification #classification of specimens
Mcluster.phenodata$uncertainty #the uncertainty of the classification
attributes(Mcluster.phenodata)
#Mclust VVE (ellipsoidal, equal orientation) model with 6 components: 
#log-likelihood   n  df       BIC       ICL
#261.2505         307 215 -708.7712 -737.2176
#Clustering table:
 # 1  2  3  4  5  6 
#26 79 22 58 78 44
#graph morphogroups according to the best model
plot(Mcluster.phenodata, what="classification", dimens=c(1,2))
#graph empirical support for different models
plot(Mcluster.phenodata, what="BIC")
#graph empirical support for the best models for each number of morphogroups
BIC.Best.Model.Per.G <- apply(Mcluster.phenodata$BIC, 1, max, na.rm=T)
max.BIC <- max(BIC.Best.Model.Per.G)
#par(mar=c(5,4,4,2)+0.1) #default
par(mar=c(5,6,4,2))
plot(1:10, max.BIC-BIC.Best.Model.Per.G[1:10], type="n", bty="n", xlim=c(1,10), ylim=c(1300,0), yaxt="n", xaxt="n",
     xlab="Number of morphological groups", ylab=expression(paste("Empirical support (",Delta, "BIC)", sep="")), 
     main="", cex.axis=1.5, cex.lab=1.5, cex.main=1.5)
points(1:10, max.BIC-BIC.Best.Model.Per.G[1:10], cex=2, pch=20, col="black", lwd=1)
#show the best Mclust models
#add abscissa
axis(1, at=c(1,seq(2,10,1)), labels=T, tcl=-0.5, cex.axis=1.5)
axis(2, at=seq(1500,0,-50), tcl=-0.7, cex.axis=1.5)
abline(v=Mcluster.phenodata$G, lty=3) #to identify the model with the most support

#"PCR"
mclust.options(hcUse="PCR")
Mcluster.phenodata <- Mclust(data.for.GMM)
#examine results
Mcluster.phenodata
summary(Mcluster.phenodata)
names(Mcluster.phenodata$classification)#the specimens included in the analysis
Mcluster.phenodata$classification #classification of specimens
Mcluster.phenodata$uncertainty #the uncertainty of the classification
attributes(Mcluster.phenodata)
#Mclust VVE (ellipsoidal, equal orientation) model with 6 components: 
# log-likelihood   n  df       BIC       ICL
#261.2505   307      215 -708.7712 -737.2176
#Clustering table:
 # 1  2  3  4  5  6 
#26 79 22 58 78 44
#graph morphogroups according to the best model
plot(Mcluster.phenodata, what="classification", dimens=c(1,2))
#graph empirical support for different models
plot(Mcluster.phenodata, what="BIC")
#graph empirical support for the best models for each number of morphogroups
BIC.Best.Model.Per.G <- apply(Mcluster.phenodata$BIC, 1, max, na.rm=T)
max.BIC <- max(BIC.Best.Model.Per.G)
#par(mar=c(5,4,4,2)+0.1) #default
par(mar=c(5,6,4,2))
plot(1:10, max.BIC-BIC.Best.Model.Per.G[1:10], type="n", bty="n", xlim=c(1,10), ylim=c(1300,0), yaxt="n", xaxt="n",
     xlab="Number of morphological groups", ylab=expression(paste("Empirical support (",Delta, "BIC)", sep="")), 
     main="", cex.axis=1.5, cex.lab=1.5, cex.main=1.5)
points(1:10, max.BIC-BIC.Best.Model.Per.G[1:10], cex=2, pch=20, col="black", lwd=1)
#show the best Mclust models
#add abscissa
axis(1, at=c(1,seq(2,10,1)), labels=T, tcl=-0.5, cex.axis=1.5)
axis(2, at=seq(1500,0,-50), tcl=-0.7, cex.axis=1.5)
abline(v=Mcluster.phenodata$G, lty=3) #to identify the model with the most support

#"SDV"
mclust.options(hcUse="SVD")
Mcluster.phenodata <- Mclust(data.for.GMM)
#examine results
Mcluster.phenodata
summary(Mcluster.phenodata)
names(Mcluster.phenodata$classification)#the specimens included in the analysis
Mcluster.phenodata$classification #classification of specimens
Mcluster.phenodata$uncertainty #the uncertainty of the classification
attributes(Mcluster.phenodata)
#Mclust VVE (ellipsoidal, equal orientation) model with 6 components: 
#log-likelihood   n  df       BIC       ICL
#261.2505       307 215 -708.7712 -737.2176
#Clustering table:
#1  2  3  4  5  6 
#26 79 22 58 78 44
#graph morphogroups according to the best model
plot(Mcluster.phenodata, what="classification", dimens=c(1,2))
#graph empirical support for different models
plot(Mcluster.phenodata, what="BIC")
#graph empirical support for the best models for each number of morphogroups
BIC.Best.Model.Per.G <- apply(Mcluster.phenodata$BIC, 1, max, na.rm=T)
max.BIC <- max(BIC.Best.Model.Per.G)
#par(mar=c(5,4,4,2)+0.1) #default
par(mar=c(5,6,4,2))
plot(1:10, max.BIC-BIC.Best.Model.Per.G[1:10], type="n", bty="n", xlim=c(1,10), ylim=c(1300,0), yaxt="n", xaxt="n",
     xlab="Number of morphological groups", ylab=expression(paste("Empirical support (",Delta, "BIC)", sep="")), 
     main="", cex.axis=1.5, cex.lab=1.5, cex.main=1.5)
points(1:10, max.BIC-BIC.Best.Model.Per.G[1:10], cex=2, pch=20, col="black", lwd=1)
#show the best Mclust models
#add abscissa
axis(1, at=c(1,seq(2,10,1)), labels=T, tcl=-0.5, cex.axis=1.5)
axis(2, at=seq(1500,0,-50), tcl=-0.7, cex.axis=1.5)
abline(v=Mcluster.phenodata$G, lty=3) #to identify the model with the most support


###################################################################################################################
###################################################################################################################
# 5) Examine phenotypic groups in the best normal mixture model.
###################################################################################################################
###################################################################################################################

###################################################################################################################
# 5.1) Examine the parameters of the multivariate normal distributions defining each phenotypic group according
# to the best normal mixture model. 

Mcluster.phenodata$parameters

###################################################################################################################
# 5.2) Examine the cross tabulation of variables in the original data frame (phenodata) and particular phenotypic
# groups in the best normal mixture model.

#select a morphogroup in the normal mixture model
mg.nmm <- 6
#select a column in original data frame (phenodata)
col.phenodata <- 4 
table(phenodata[as.numeric(names(Mcluster.phenodata$classification))[Mcluster.phenodata$classification==mg.nmm],col.phenodata])

###################################################################################################################
# 5.3) Graph empirical support for the best models for each number of phenotypic groups.

BIC.Best.Model.Per.G <- apply(Mcluster.phenodata$BIC, 1, max, na.rm=T)
max.BIC <- max(BIC.Best.Model.Per.G)
#par(mar=c(5,4,4,2)+0.1) #default
par(mar=c(5,6,4,2))
plot(1:10, max.BIC-BIC.Best.Model.Per.G[1:10], type="n", bty="n", xlim=c(1,10), ylim=c(1300,0), yaxt="n", xaxt="n",
     xlab="Number of morphological groups", ylab=expression(paste("Empirical support (",Delta, "BIC)", sep="")), 
     main="", cex.axis=1.5, cex.lab=1.5, cex.main=1.5)
points(1:10, max.BIC-BIC.Best.Model.Per.G[1:10], cex=2, pch=20, col="black", lwd=1)
#show the best Mclust models
#add abscissa
axis(1, at=c(1,seq(2,10,1)), labels=T, tcl=-0.5, cex.axis=1.5)
axis(2, at=seq(1500,0,-50), tcl=-0.7, cex.axis=1.5)
abline(v=Mcluster.phenodata$G, lty=3) #add a line showing the model with highest empirical support

###################################################################################################################
# 5.4) Plot phenotypic groups in the best normal mixture model. 

par (mfcol=c(2,2),mar =c(2,2,2,2),oma=c(0.9,1.4,1,0.6),mai=c(0.7,0.7,0.2,0.2))
plot(Mcluster.phenodata, what=c("classification"), dimens=c(1,2), main="")
#plot(Mcluster.phenodata, what=c("uncertainty"), dimens=c(1,2), main="")
plot(Mcluster.phenodata, what=c("classification"), dimens=c(1,3), main="")
#plot(Mcluster.phenodata, what=c("uncertainty"), dimens=c(1,3), main="")
plot(Mcluster.phenodata, what=c("classification"), dimens=c(3,2), main="")
#plot(Mcluster.phenodata, what=c("uncertainty"), dimens=c(3,2), main="")
#plot legend
plot(c(0,0), type="n", bty="n", xaxt="n", yaxt="n",xlab="",ylab="" )
legend("center", paste("Morphological group", 1:6),
	col=c("dodgerblue2","red3","green3","slateblue","darkorange","skyblue1","violetred4","forestgreen","steelblue4"),
	pch=c(16,0,17,3,15,4,1,8,2), pt.lwd=1, pt.cex=1, cex=1.3, bty="o")

###################################################################################################################
# 5.5) Plot loadings for different phenotypic traits on the principal component space. 

par(mfcol=c(2,2), mar=c(5,5,2,2)+0.1)
#plot loadings PC1 and PC2
plot(0,0, xlim=c(-0.76,0.76), ylim=c(-0.76,0.76), asp=1, type="n", bty="n",
     xlab = "Loadings PC1", ylab = "Loadings PC2",
     cex.axis=1, cex.lab=1)
for (i in 1:13)
	{
	arrows(0, 0, x1 = mean.phenodata.selected.log.pca$rotation[,1][i], y1 = mean.phenodata.selected.log.pca$rotation[,2][i], 
	length = 0.1, angle = 20, code = 2, col = "black", lwd=1)
	}
#plot circle for equal contribution of traits.
x <- seq(-(2/13)^0.5, (2/13)^0.5, 1e-4) 
equal_contribution<-((2/13)-x^2)^0.5
points(x, equal_contribution, type="l", lty=1, col="gray70")
points(x, -equal_contribution, type="l", lty=1, col="gray70")
for (i in 1:13)
	{
	text(mean.phenodata.selected.log.pca$rotation[,1][i]*1.1, mean.phenodata.selected.log.pca$rotation[,2][i]*1.1, i, col="cyan")
	}

#plot loadings PC1 and PC3
plot(0,0, xlim=c(-0.76,0.76), ylim=c(-0.76,0.76), asp=1, type="n", bty="n",
     xlab="Loadings PC1", ylab="Loadings PC3", cex.axis=1, cex.lab=1)
for (i in 1:13)
	{
	arrows(0, 0, x1 = mean.phenodata.selected.log.pca$rotation[,1][i], y1 = mean.phenodata.selected.log.pca$rotation[,3][i], 
	length = 0.1, angle = 20, code = 2, col = "black", lwd=1)
	}
#plot circle for equal contribution of traits.
x <- seq(-(2/13)^0.5, (2/13)^0.5, 1e-4) 
equal_contribution<-((2/13)-x^2)^0.5
points(x, equal_contribution, type="l", lty=1, col="gray70")
points(x, -equal_contribution, type="l", lty=1, col="gray70")
for (i in 1:13)
	{
	text(mean.phenodata.selected.log.pca$rotation[,1][i]*1.1, mean.phenodata.selected.log.pca$rotation[,3][i]*1.1, i, col="cyan3")
	}

#plot loadings PC3 and PC2
plot(0,0, xlim=c(-0.76,0.76), ylim=c(-0.76,0.76), asp=1, type="n", bty="n",
     xlab="Loadings PC3", ylab="Loadings PC2", cex.axis=1, cex.lab=1)
for (i in 1:13)
	{
	arrows(0, 0, x1 = mean.phenodata.selected.log.pca$rotation[,3][i], y1 = mean.phenodata.selected.log.pca$rotation[,2][i], 
	length = 0.1, angle = 20, code = 2, col = "black", lwd=1)
	}
#plot circle for equal contribution of traits.
x <- seq(-(2/13)^0.5, (2/13)^0.5, 1e-4) 
equal_contribution<-((2/13)-x^2)^0.5
points(x, equal_contribution, type="l", lty=1, col="gray70")
points(x, -equal_contribution, type="l", lty=1, col="gray70")
for (i in 1:13)
	{
	text(mean.phenodata.selected.log.pca$rotation[,3][i]*1.1, mean.phenodata.selected.log.pca$rotation[,2][i]*1.1, i)
	}

rownames(mean.phenodata.selected.log.pca$rotation)
trait.names <- c("Log (lamina length)", "Log (lamina width)", "Log (synflorescence axis length)",
	"Log (synflorescence axis perimeter)", "Log (sterile leaf pairs per synflorescence + 1)",
	"Log (capitula per synflorescence)", "Log (terminal cyme peduncle length)",
	"Log (capitulum diameter)", "Log (sterile phyllary length)", "Log (sterile phyllary width)",
	"Log (ray flower corolla length)", "Log (disc flower corolla length)",
	"Log (disc flower tube length)") 
#par(mar=c(5,4,4,2)+0.1) #default
par (mfrow=c(1,1), mar=c(5,4,4,2)+0.1)
plot(c(-0.1,1), c(-0.1,1), type="n", bty="n", xaxt="n", yaxt="n", xlab="", ylab="")
legend(x=0, y=1, paste(1:13, ". ", trait.names, sep=""), col=("black"), pt.lwd="0", cex=1)
